# Effect of Perinatal Ampicillin or Amoxicillin/Clavulanate Exposure on Maternal and Infant Gut Microbiome, Metabolome, and Infant Responses to the 20-valent Pneumococcal Conjugate Vaccine

**DOI:** 10.64898/2025.12.08.692990

**Authors:** Emi Suzuki, Victoria Deleray, Jasmine Zemlin, Armin Kousha, Hannah Nonoguchi, Daniel Sun, Chih-Ming Tsai, Simone Zuffa, Kine Eide Kvitne, Pieter C. Dorrestein, Shirley M. Tsunoda, Victor Nizet, George Y. Liu, Fatemeh Askarian

## Abstract

Emerging studies suggest that antibiotics can disrupt the gut microbiome and alter vaccine-induced immune responses, but the specific consequences of early-life exposure on neonatal immune development remains poorly understood. Here, we examined how two antibiotics frequently used in perinatal care, broad-spectrum ampicillin (AMP) and the extended-spectrum combination amoxicillin/clavulanate (AMOX/CLAV), administered during gestation and lactation, influence neonatal gut microbiome composition, fecal metabolome profiles, and responses to the 20-valent pneumococcal conjugate vaccine (PCV20). Maternal treatment with AMOX/CLAV, but not AMP, significantly reduced PCV-specific IgG titers at 4-and 6-weeks post-prime immunization compared to untreated controls. Exclusive exposure to AMOX/CLAV also impaired neutrophil-mediated opsonophagocytic killing, indicating diminished antibody functionality. These effects were transient, with immune parameters normalizing by week 8 post-prime immunization. Metabolomic and microbiome profiling revealed that maternal AMP and AMOX/CLAV differentially perturbed specific metabolite classes including bile acids, *N*-acyl lipids, and indole-derivatives, as well as key commensal taxa including Bacteroidales and Coriobacteriales within the gut microbiota. Together, these findings reveal a previously underappreciated maternal-offspring route of antibiotic influence that transiently disrupts neonatal vaccine responsiveness through microbiome and metabolome alterations. These results highlight maternal antibiotic exposure as a modifiable factor shaping early-life immunity.

## INTRODUCTION

Prophylactic administration of β-lactam antibiotics, such as ampicillin and penicillin is a standard obstetric practice following prenatal identification of maternal group B *Streptococcus* (GBS) colonization. This intrapartum intervention has been robustly linked to a significant reduction in early-onset GBS disease in neonates, underscoring its critical role in contemporary perinatal infection control.^1,2^ It is estimated that up to 40% of neonates in North America are indirectly exposed to intrapartum antibiotics, regardless of delivery mode (vaginal vs. caesarean).^3,4^ Although breastfeeding supports microbial and immune development, even a single course of intrapartum antibiotic course can alter the infant gut microbiota, with changes persisting for up to three months after birth.^3^

The early postnatal period represents a critical window for immune system priming, during which gut microbial cues profoundly influence long-term immune trajectories.^5,6,7,8^ Disruption of this process, particularly from early-life antibiotic exposure, has been associated with adverse outcomes, including the emergence of disease phenotypes in adulthood (reviewed in ^9^), impaired vaccine responsiveness in murine models^10^ and human infants,^11–14^ increased neonatal susceptibility to sepsis^15^ and the development of antibiotic resistance. Alarmingly, global estimates indicate that 30-50% of pregnant or lactating individuals are prescribed antibiotics,^16,17^ raising concerns about antibiotic resistance,^18^ disruption of neonatal microbiota, and consequent immune dysregulation.

Beyond acute effects, early antibiotic exposure has been linked, epidemiologically or experimentally, to an elevated risk of immune-mediated disorders such as atopic dermatitis, asthma, type 1 diabetes, metabolic syndrome, obesity, eosinophilic esophagitis, and inflammatory bowel disease (IBD).^19–24,25,26,27^ Paradoxically, under certain conditions defined by timing, dose, antibiotic class, and infant health background,^28,29^ antibiotic exposure may instead confer protection, as reported for atopic dermatitis in infants^30^ and IBD in a murine model.^30,31^ These seemingly contradictory outcomes underscore the complexity and context dependence of host-microbiome-immune system interactions and highlight the need to elucidate the immunological consequences of antibiotic-induced microbiome perturbation during this formative window.

Antibiotic exposure during infancy is consistently associated with reduced *Bifidobacterium* abundance,^32–37^ a dominant early-life taxon linked to enhanced vaccine efficacy.^11,38^ Retrospective studies further associate early antibiotic exposure with attenuated antibody responses to routine childhood immunizations,^12^ with prolonged broad-spectrum antibiotic courses causing greater suppression of vaccine-specific antibody titers than shorter narrow-spectrum regimens.^12^ To dissect the impact of antibiotic exposure on vaccine-induced immunity in offspring, we employed a murine model of early-life antibiotic exposure to examine the immunological consequences of maternal perinatal treatment with ampicillin (AMP), the most commonly prescribed agent in peripartum prophylaxis protocols, or the extended-spectrum combination amoxicillin/clavulanate (AMOX/CLAV). We evaluated neonatal responses to the 20-valent pneumococcal conjugate vaccine (PCV20), alongside detailed gut microbiome and fecal metabolomic profiling. Our integrated analyses uncovered distinct alterations in microbial communities and metabolic outputs that correlated with vaccine-induced humoral protection. These findings highlight the microbiome’s pivotal role in modulating vaccine efficacy and suggest that the breadth of antibiotic coverage during early life may differentially shape immune development through microbiota-metabolite-host interactions.

## MATERIALS AND METHODS

### Ethics declarations

All animal experiments were conducted in compliance with ethical guidelines for animal research, following Institutional Animal Care and Use Committee regulations and approved under UC San Diego IRB protocol S00227M and S18200.

### Murine model of antibiotic administration during gestation and postpartum

C57BL/6J (The Jackson Laboratory, JAX:000664) mice were purchased and were housed throughout the experiment in filter-top cages with free access to commercial chow diet (2020X Teklad Global Soy Protein-Free Extruded Rodent Diets, inotiv/ENVIGO) and water *ad libitum* under conditions of regulated ambient temperature (20-22 °C) and relative humidity (30-70%), with a 12 h light/12h dark cycle. Timed pregnancies were established by pairing 8-to 10-week-old male and female mice (1:1) for 24 hours, after which females were single housed. Pregnancy was confirmed on gestational day 14 (G14) by visible abdominal enlargement and abdominal palpation. Treatment commenced on embryonic day 16 (E16) and pregnant female mice (n= 18 dams; 1 mouse per cage) and mice were randomized into the designated experimental treatments including mock (phosphate-buffered saline: PBS; Sigma Aldrich), ampicillin (AMP) or amoxicillin/clavulanic acid (AMOX/CLAV). Treatment continued through day 7 (D7; Wk1) postpartum, targeting critical periods of gestation and early postpartum advancement. Mice in the antibiotic treatment arm were exposed to AMP (1 mg/ml; 150-300 mg/kg/day assuming dams consume 4-8 ml water per day) or AMOX/CLAV (0.5 mg/ml; 75-150 mg/kg/day), which were dissolved in drinking water and provided continuously to the dams. Water was replaced every 48 hours throughout the treatment period to maintain consistent antibiotic exposure. Mice gave birth on gestation day 19 (G19), and the number of infants in each treatment group was reported. Specifically, the number of pups from PBS-, AMP-and AMOX/CLAV-treated dams were 38 (from 6 timed-pregnant dams), 29 (from 5 timed-pregnant dams), and 45 (from 7 timed-pregnant dams), respectively. For microbiome and metabolome analyses, fecal samples were collected from dams at week 0 (prior initiation of oral antibiotic treatment), and subsequently at weeks 1, 2, 3 and 4 postpartum. Fecal samples from pups were collected at weeks 2, 3, and 4 post-birth. Fecal pellets were carefully harvested using sterile forceps and immediately frozen at-80 °C until analysis. To assess the transfer of AMP or AMOX/CLAV to the pups, peripheral blood was collected via submandibular bleed from both dams and pups at week 1, following antibiotic exposure of the dams. Murine sera were isolated via centrifugation and stored at-80 °C until analysis.

### Post-natal immunization with 20-valent pneumococcal conjugate vaccine (PCV20)

All pups were randomly assigned (per treatment) to receive intraperitoneal immunization with 100 µL of the PCV20 (AMP: n = 16, AMOX/CLAV: n = 29, Mock: n = 20 (Pfizer), diluted 1:2 in sterile PBS, or a sham immunization with 100 µL of PBS (AMP: n = 13, AMOX/CLAV: n = 16, Mock: n = 18) on postnatal days 14 (D14 or Wk2) and 28 (D28 or Wk4). Peripheral blood was randomly collected via submandibular bleed from PCV20-(AMP: n = 6, AMOX/CLAV: n = 6, Mock: n = 6) and sham-immunized mice (AMP: n = 5, AMOX/CLAV: n = 6, Mock: n = 6) at weeks 2, 4, 6, and 8 following the initial immunization (prime). Murine sera were isolated via centrifugation and stored at-80 °C until analysis.

### Quantification of PCV20-specific antibodies in infant murine sera

The levels of PCV20-specific antibodies, including IgG total, IgG1, IgG2b, and IgA were quantified in serum from infant pups using enzyme-linked immunosorbent assay (ELISA). Briefly, high-binding microtiter plates were coated with 100 µL of diluted PCV20 in PBS (1:100) and incubated overnight at 4°C. The wells were then blocked with 1% (w/v) bovine serum albumin (BSA) for 2 hours at room temperature (RT), followed by washing with PBS supplemented with 0.05% (v/v) Tween 20 (Sigma Aldrich). Serially diluted sera in PBS (100 μL) were added to the wells and incubated for 2 hours at RT. Bound antibodies were detected using HRP-conjugated goat anti-mouse IgG (BioLegend # 405306, dilution: 1: 5,000), biotin-conjugated anti-mouse IgA (BioLegend # 407003, dilution: 1:5,000), biotin-conjugated anti-mouse IgG1 (BioLegend # 406603, dilution: 1:5,000), and biotin-conjugated anti-mouse IgG2b (BioLegend # 406703, dilution: 1:5,000). Detection was performed with secondary HRP-avidin antibodies at a 1:1,000 using TMB substrate kit (BD OptEIA). The plates were read by plate reader at 450 nm, with background subtraction at 570 nm for correction.

### Opsonophagocytosis assay for functional antibody evaluation against *Streptococcus pneumoniae*

Opsonophagocytic killing assays were conducted using primary murine neutrophils, as described previously with modifications^39^ using *S. pneumoniae* (SPN) strain TIGR4. Mouse neutrophils were isolated from bone marrow by MojoSort Mouse Neutrophil Isolation Kit, according to the manufacturer’s instruction (Biolegend). An overnight culture of SPN was grown at 37°C in Todd Hewitt broth supplemented with 2 % (w/v) yeast extract (THY), and frozen glycerol stocks were made. The frozen stock was thawed, and bacteria were washed with 1x THY and resuspended with THY. SPN was opsonized with mouse serum for 20 minutes at 37°C with continuous agitations. Opsonized SPN was incubated with freshly isolated mouse neutrophils (2 x 10^5^) at a multiplicity of infection (MOI) of 1:0.01 (neutrophils: SPN) in the presence of 2% (v/v) mouse serum. Following a 1-hour incubation at 37°C, serial dilutions were plated on blood agar plates (Hardy Diagnostics, A10BX) for enumeration of surviving colony-forming units (CFU).

### Metabolomics

A single fecal pellet per sample was extracted with 800 μL of 50% LC-MS grade methanol and homogenized by bead beating with a 5mm bead at 25 Hz for 5 minutes, followed by a 30-minute incubation at 4 °C. The mixture was centrifuged at 15,000g for 5 minutes in order to pellet the precipitate and solid material. The supernatant was transferred to a new 96-well plate then vacuum concentrated via centrifugal lyophilization (Labconco Centrivap) until ready for LC/MS analysis. The resulting supernatant was resuspended in 400 μL 50% LC/MS grade methanol in 96 well plates with 1 μL sulfadimethoxine as an internal standard. The fecal extracts were analyzed on a Thermo™ QExactive™ mass spectrometer coupled to a Vanquish Ultra-High-Performance Liquid Chromatography system (ThermoFisher). Chromatographic separation was performed on a reverse-phase HPLC C18 column 2.1mm x 120mm using 0.1% formic acid in water (A) and 0.1% formic acid in acetonitrile (B) as the mobile phase. The chromatography ran in a 12-minute gradient: 0-1 min 1% B, 1-7.5 min 1-99% B, 7.5-9.3 min 99% B, 9.3-9.5 min 99-1% B, 9.5-11 min 1% B. The column compartment was kept at 40 °C and the sample injection volume was 3 μL with a flow rate of 0.5 mL per minute. Detection was performed in electrospray ionization positive mode using a data-dependent acquisition method with the following parameters: nebulizer gas pressure at 2 bar, spray voltage was set to 3.5 kV, and sheath gas flow rate set to 54. Full MS1 scans of *m/z* 100-1300 (resolution=35,000) were followed by MS2 acquisition of the five most intense precursor ions per scan (resolution=17,500) at 30 eV collision energy. The raw files were converted to.mzML format using Proteowizard^40^ MZConvert software for data processing and analysis in downstream softwares. LC-MS/MS data for these fecal extractions are available in the MassIVE data repository under accession number <u>MSV000097137</u>. The.mzML files were analyzed in MZmine 3.9.2^41^ under the following parameters: MS1 noise factor of lowest signal 5, MS2 noise factor of lowest signal 2.5 with 1000 minimum intensity and 5000 minimum feature height with at least 4 scans per feature. The *m/z* tolerance was 0.02 or 10 ppm and the retention time tolerance was 0.1 minutes. The isotopic peaks finder, isotope grouper, and metaCorrelate modules were applied to identify multiple ion forms per molecule. The export to GNPS function exported a feature table with peak area abundances and a spectral text file. Feature-based molecular networking (FBMN) in GNPS2^42^ was applied to the filtered features with *m/z* tolerance 0.02, minimum cosine similarity score 0.7, and minimum matching peaks of 4. Library matching to “All GNPS” library and the propagated bile acid candidate library^43^ used the same tolerances. Further inspection of bile acid spectra included using previously developed MassQL queries^44^ for validation of spectra and knowledge of core structures, and manual investigation of the propagations of validated spectra within the feature based molecular network. Acylated carnitines were annotated using an in-house spectral library and manual investigation of the propagations of validated spectra within the network by a diagnostic fragment ion of 84.08 Da. The CMMC enrichment workflow in GNPS2 annotated microbial derived *N*-acyl lipids^45^. The peak area data was processed in RStudio version 4.4.0 to remove features with high signal in blank extractions (sample/blank > 5) and internal standards. Internal standards were used to evaluate data quality. Package vegan 2.6.10 was used for rclr transformation and mixOmics 6.28.0 was used for principal component analysis (PCA) and partial least squares discriminant analysis (PLS-DA). Centroid separation in PCA was evaluated with Permutational multivariate analysis of variance (PERMANOVA). PLSDA model performance was assessed using 4-fold cross-validation and 100 repeats. PLS-DA, dotplot with heatmap, and boxplots visualizations were created using ggplot2 3.5.1. Variable Importance Projection (VIP) scores were calculated and extracted from the PLSDA model. Features with VIP scores > 1 that also had annotations from FBMN were retained and used for further analysis. Univariate statistical tests were performed using Wilcoxon test followed by Benjamini-Hochberg (BH) correction.

### Microbiome analysis

The UC San Diego Microbiome Core performed nucleic acid extractions utilizing previously published protocols.^46^ Briefly, samples were purified using the MagMAX Microbiome Ultra Nucleic Acid Isolation Kit (Thermo Fisher Scientific, USA) and automated on KingFisher Flex robots (Thermo Fisher Scientific, USA). Blank controls and mock communities (Zymo Research Corporation, USA) were included per extraction plate, which were carried through all downstream processing steps. DNA was quantified using a PicoGreen fluorescence assay (Thermo Fisher Scientific, USA) and 16S rRNA gene amplification was performed according to the Earth Microbiome Project protocol. ^47^ Briefly, Illumina primers with unique forward primer barcodes were used to amplify the V4 region of the 16S rRNA gene (515fB-806r, ^48^). Amplification was performed in a miniaturized volume,^49^ with single reactions per sample.^50^ Libraries were subsequently pooled, then sequencing was carried out at the UC San Diego Institute for Genomic Medicine on the Illumina MiSeq sequencing platform with Reagent Kit v2 and paired-end 150 bp cycles. Raw data was imported in Qiita (<u>Qiita #15920</u>)^51^ and processed using Deblur^52^ default workflow. The resulting BIOM table was processed in Qiime2-amplicon-2024.10^53^. Greengenes2 was used for phylogeny and taxonomy^54^. Samples with less than 10,000 reads were removed, and ASVs present in less than 10% of samples were removed. To control for sequencing effort, the dataset was rarefied to 10,000 reads for the Shannon diversity analysis. The feature tables were converted to. tsv format from Qiime2 for import into RStudio version 4.4.0 for downstream analysis. Package vegan 2.6.10 was used for rclr transformation and mixomics 6.28.0 was used for PCA. Centroid separation was evaluated with PERMANOVA. Shannon entropy was calculated using the Qiime2 environment and analyzed in RStudio. Statistical significance of alpha diversity was assessed using a linear mixed-effects model for repeated measures. ALDEx2^55^ was used for differential abundance analysis of antibiotic exposure against mock exposure and BH adjusted *P* < 0.05 were considered significant. The integrative analysis of the rclr transformed microbiome feature table and the rclr-transformed metabolomic feature table was performed using Data Integration Analysis for Biomarker discovery using Latent Components (DIABLO).^56^ To maximize covariance between groups, the DIABLO model was tuned to keep top features, then performance measured using a leave-one-out (loo) cross validation and 500 permutations. The chosen correlating features were visualized in a circos plot where a correlation >0.7 were retained.

### Quantification and statistical analysis

Data were processed and visualized using GraphPad Prism (version 10). Results are expressed as the mean ± standard error of the mean (SEM), as indicated in the figure legends. Statistical significance (*P* < 0.05) was assessed using one-way ANOVA. Further details, including the specific statistical test used, corresponding *P*-value, sample size (number of mice), and biological replicates, can be found in the figure legend. Statistical analyses for metabolomics and microbiome data are described in the respective section.

## RESULTS

### Peripartum exposure to AMOX/CLAV transiently attenuates PCV20-specific IgG and neutrophil-mediated opsonophagocytic killing in offspring

Dams were treated with oral AMP or AMOX/CLAV from late gestation (E16) to the end of first postpartum week (Wk1) at standard weight-based dosing (total 10 days). Pups received PCV20 at Wk2 (prime) and Wk4 (boost), and PCV20-specific antibody responses were assessed as outlined in Figure 1A. Maternal AMOX/CLAV exposure significantly reduced total PCV20-specific IgG titers at 4-and 6-weeks post-prime compared with pups from AMP-treated or untreated dams, an effect that was resolved by week 8 post-prime (Fig. 1B). Subclass analysis demonstrated statistically significantly lower IgG1 and IgG2b titers in AMOX/CLAV-exposed pups at week 4 (Fig. 1D-E), whereas PCV20-specific IgA remained uniformly low across groups (Fig. 1C).

**Figure 1.**
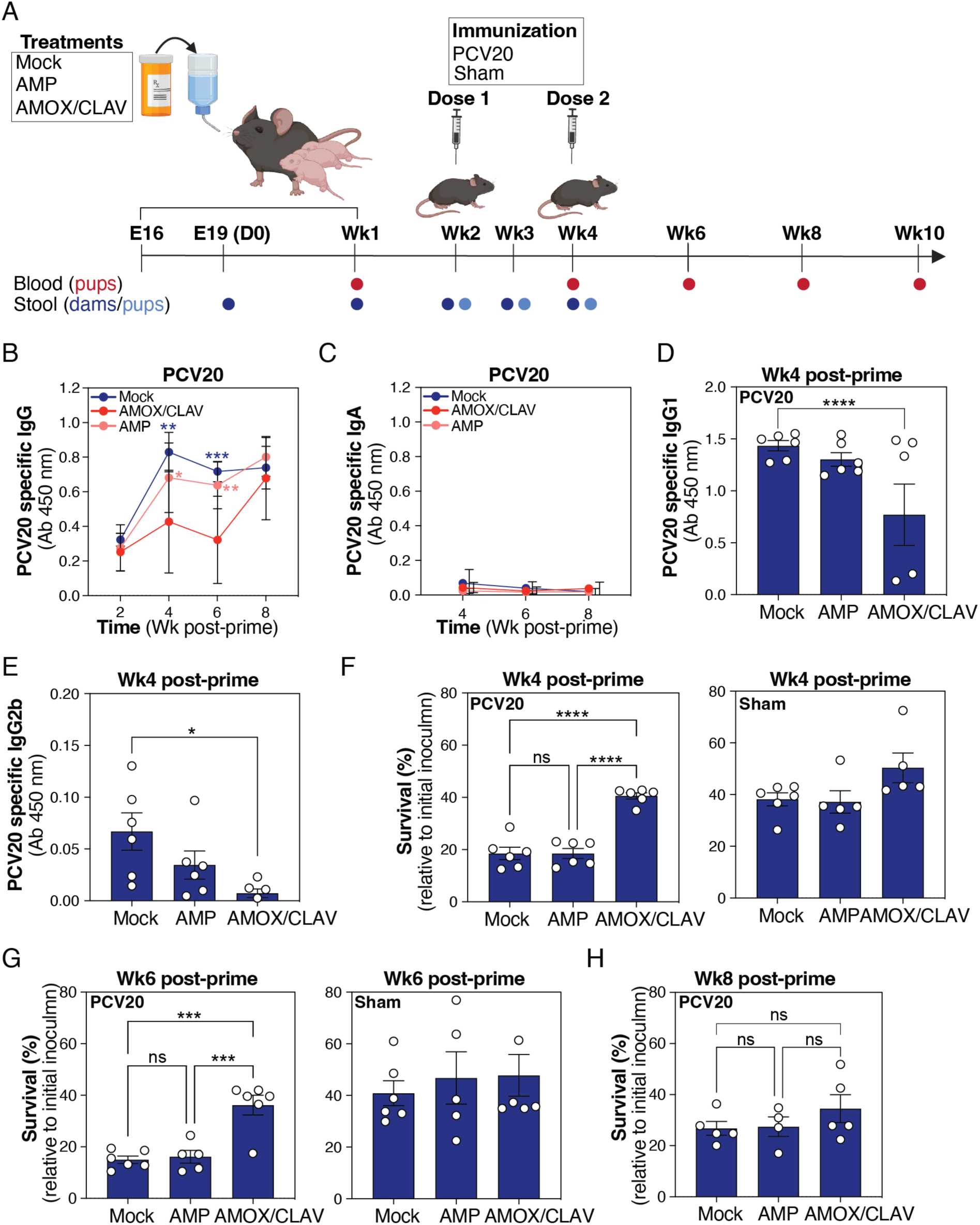
Peripartum exposure of dams to AMOX/CLAV and their immunological response to PCV20 immunization. **(A)** Schematic overview of experimental design and sampling. Dams were either left untreated (mock), administered oral ampicillin (AMP) or oral amoxicillin/clavulanate (AMOX/CLAV) beginning in late prepartum (E16) and continuing through the end of first postpartum week (Wk1). Pups received either the 20-valent pneumococcal conjugate vaccine (PCV20) or a sham immunization. Blood samples (red) and stool samples (moms: dark blue; pups: light blue) were collected at the defined timepoints. Experimental details are provided in the materials and methods section. The figure was initially created with BioRender and subsequently refined in Adobe Illustrator. **(B)** Total serum anti-PCV20 IgG levels were measured following PCV20 immunization at the dose of 100 µL/mouse (PCV20 diluted 1:2 in PBS). Serum samples were collected at weeks 2, 4, 6, and 8 post-prime as described in A. The non-immunized (sham) control is shown in Fig. S1A. Data are plotted as the mean ± SEM, representing 4 - 6 mice per group, and were analyzed by two-way ANOVA (Tukey’s multiple comparisons). Wk4: Mock vs AMOX/CLAV: *P* = 0.0005, AMP vs AMOX/CLAV: *P* = 0.0355; Wk6: Mock vs AMOX/CLAV: *P* = 0.0006, AMP vs AMOX/CLAV: *P* = 0.0068. **(C)** Total serum anti-PCV20 IgA levels were measured following PCV20 immunization as described in B. Serum samples were collected at weeks 4, 6, and 8 post-prime as described in A. The non-immunized (sham) control is shown in Fig. S1B. Data are plotted as the mean ± SEM, representing 5 - 6 mice per group, and were analyzed by two-way ANOVA (Tukey’s multiple comparisons). **(D)** Total serum anti-PCV20 IgG1 levels were measured following PCV20 immunization as described in B. Serum samples were collected at week 4 post-prime. The non-immunized (sham) control is shown in Fig. S1C. Data are plotted as the mean ± SEM, representing 6 mice per group, and were analyzed by one-way ANOVA (Tukey’s multiple comparisons). Mock vs AMOX/CLAV: *P* = 0.0457. **(E)** Total serum anti-PCV20 IgG2b levels were measured following PCV20 immunization as described in B. Serum samples were collected at week 4 post-prime. The non-immunized (sham) control is shown in Fig. S1D. Data are plotted as the mean ± SEM, representing 6 mice per group, and were analyzed by one-way ANOVA (Tukey’s multiple comparisons). Mock vs AMOX/CLAV: *P* = 0.0157. **(F)** Opsonophacytic activity of anti-PCV20 IgG was evaluated using an *in vitro* opsonophagocytic killing assay. SPN was incubated (60 min) with freshly isolated murine neutrophils (MOI = 0.01) in the presence of 10% serum obtained from PCV20-immunized (left panel) or sham-immunized (right panel) mice at week 4 post-prime. Data are plotted as the mean ± SEM, representing 5 - 6 mice per group, and were analyzed by one-way ANOVA (Tukey’s multiple comparisons). PCV20: Mock vs AMOX/CLAV: *P* < 0.0001, AMP vs AMOX/CLAV: *P* < 0.0001. **(G)** Opsonophagocytic activity of anti-PCV20 IgG was evaluated using an *in vitro* opsonophagocytic killing assay. SPN was incubated (60 min) with freshly isolated murine neutrophils (MOI = 0.01) in the presence of 10% serum obtained from PCV20-immunized (left panel) or sham-immunized (right panel) mice at week 6 post-prime. Data are plotted as the mean ± SEM, representing 5 - 6 mice per group, and were analyzed by one-way ANOVA (Tukey’s multiple comparisons). PCV20: Mock vs AMOX/CLAV: *P* = 0.0002, AMP vs AMOX/CLAV: *P* = 0.0006. **(H)** Opsonophagocytic activity of anti-PCV20 IgG was evaluated using an *in vitro* opsonophagocytic killing assay. SPN was incubated (60 min) with freshly isolated murine neutrophils (MOI = 0.01) in the presence of 10% serum obtained from PCV20-immunized mice at week 8 post-prime. Data are plotted as the mean ± SEM, representing 4 - 5 mice per group, and were analyzed by one-way ANOVA (Tukey’s multiple comparison).

Functional antibody activity, measured by neutrophil opsonophagocytic killing (OPK), was likewise reduced in AMOX/CLAV-exposed pups at weeks 4 and 6 post-prime (Fig. 1F-G) and recovered by week 8 (Fig. 1H).

### Maternal antibiotic exposure affects gut microbiota composition in dams and offspring

PCA of robust center log ratio (rclr)-transformed data revealed distinct clustering by treatment and time in both dams and pups (Fig. 2A), with antibiotic exposure and sampling time exerting strong effects on fecal microbiome composition (PERMANOVA, dams, treatment: *P* = 0.001, R^2^ = 0.37634, F = 81.2695, time: *P* = 0.001, R^2^ = 0.31288, F = 33.7831) (PERMANOVA, pups, treatment: *P* = 0.001, R^2^ = 0.27040, F = 25.0924, time: *P* = 0.001 R^2^ = 0.46395, F = 43.0531). No clear separation was observed between AMP and AMOX/CLAV groups. In dams, samples collected during active treatment (E19 and Wk1) clustered separately from post-treatment samples (Wk2-4), indicating a shift between exposure and recovery phases. In pups, Wk2 microbiomes clustered distinctly from Wk3-Wk4, reflecting temporal maturation following maternal treatment. Alpha diversity was reduced in antibiotic-treated dams compared with controls (PERMANOVA, *P* < 0.01; Fig. 2B). In pups, overall, Shannon diversity was similar across groups (Fig. S2A), but among immunization samples, AMOX/CLAV-exposed pups had reduced diversity compared with controls (*P* < 0.1), whereas AMP exposure did not alter diversity (*P* = 0.51)(Fig. 2B). These findings indicate that pups with reduced vaccine responses also exhibited decreased gut microbial richness.

**Figure 2.**
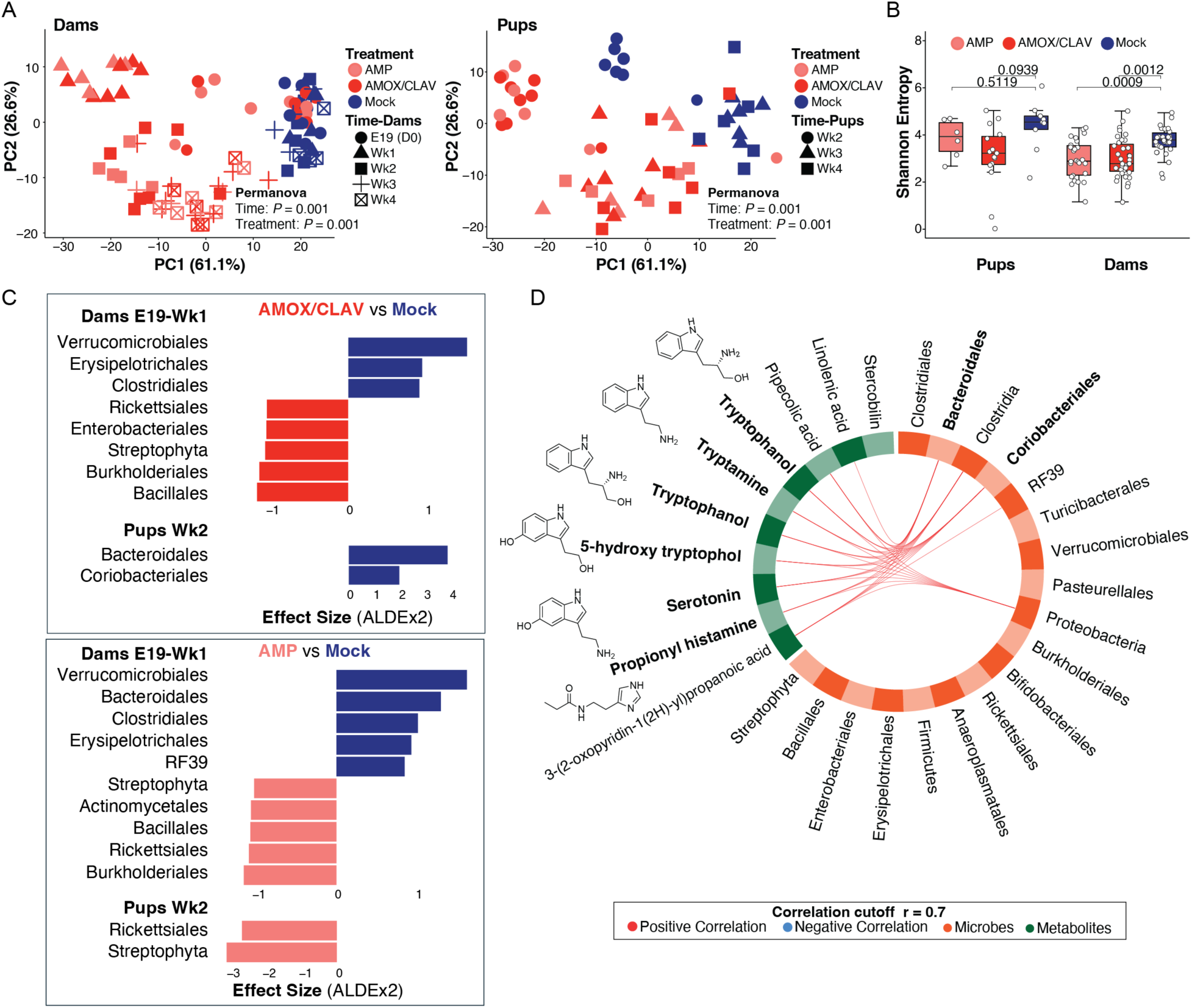
Mice exposed to antibiotics show altered microbiomes. **(A)** Principal component analysis (PCA) of rclr-transformed microbial features revealed the overall microbial separation between mice exposed to antibiotic treatment or mock. Separation was assessed using PERMANOVA, showing a significant effect of both treatment condition and time in pups and dams that were indirectly or directly exposed to antibiotics, respectively (PERMANOVA: treatment *P* = 0.001, time *P* = 0.001). **(B)** Shannon diversity was significantly reduced in dams receiving antibiotic treatments (*P* < 0.01, linear mixed-effects model (LME) with subject as random effect). Similarly, pre-and post-immunized PCV20 Wk2-4 pups that were exposed indirectly to AMOX/CLAV showed depleted Shannon diversity (*P* = 0.09, linear mixed-effects model for repeated measurements) compared to no observed difference between AMP and Mock (LME: *P* = 0.51). The boxplots represent first (lower), interquartile range (IQR), and third (upper) quartile. **(C)** Microbial orders driving differences between antibiotic treatment conditions at early time points (Dams E19-Wk1, Pups Wk2) were identified using differential abundance analysis with ALDEx2 (FDR-corrected *P* < 0.05). **(D)** Multi-omics integration of top 10 rclr-transformed annotated microbial features, collapsed at the order level, and top 10 rclr-transformed annotated metabolic features revealed strong correlations, visualized using a circos plot generated with DIABLO. Corrections greater than 0.7 were retained. Negative (none detected) and positive correlations are depicted in blue and red, respectively.

To investigate the taxonomic drivers of these differences, we performed differential abundance analysis using ALDEx2 (Fig. 2C). Samples were stratified by the timepoints and vaccination status identified in earlier analyses (Fig. 2A-B), and features annotated against Greengenes phylogeny were collapsed at the order level.

During active antibiotic treatment, AMOX/CLAV-treated dams exhibited significant depletion of several commensal orders including Verrucomicrobiales, Erysipelotrichales, and Clostridiales, with concurrent enrichment of Bacillales and Burkholderiales (Fig. 2C) (*P* < 0.05). AMP-treated dams similarly showed depletion of Verrucomicrobiales, Bacteroidales, and Clostridiales, alongside enrichment of Burkholderiales, Rickettsiales, and Bacillales. Notably, post-treatment samples contained more significantly enriched or depleted taxa than samples collected during active exposure (AMP: 19 vs. 6, AMOX/CLAV: 19 vs. 4) (Dataset 1). Although many changes overlapped between treatment groups, post-AMOX/CLAV treatment uniquely depleted Lactobacillales and enriched Pseudomonadales and Pasteurallales (Fig. S3A), whereas post-AMP treatment uniquely depleted Erysipelotrichales.

At Wk2 (prior to PCV20 immunization), pups from AMOX/CLAV-treated dams exhibited selective depletion of Bacteroidales and Coriobacteriales compared with mock controls, whereas no significantly depleted taxa were observed in AMP-exposed pups (Fig. 2C). In contrast, AMP-exposed pups showed enrichment of Streptophyta and Rickettsiales. The similarity between dam and offspring community profiles indicates maternal transmission of antibiotic-induced microbiome perturbations. By Wk3-4, AMP-exposed, PCV20-immunized pups exhibited enrichment in Turicibacterales and Clostridiales (Fig. S3A, right panel). AMOX/CLAV-exposed, PCV20-immunized pups remained uniquely depleted in Bacteroidales and Coriobacteriales, while showing enrichment in Clostridiales, Rickettsiales, Bacillales, Bifidobacteriales, Streptophyta, and Burkholderiales, (Fig. S3A, left panel) (Dataset 2). These patterns indicate that AMOX/CLAV drives more sustained depletion of commensal anaerobes associated with early immune education.

### Maternal antibiotic exposure induces coordinated and pathway-specific metabolomic shifts in offspring

Untargeted metabolomics of fecal samples was performed alongside microbiome profiling and integrated using DIABLO, a supervised multi-omics framework designed to identify features that covary across datasets. Feature selection was guided by optimization of a partial least squares (PLS) model, and metabolite-taxon associations with correlation coefficients >0.70 were retained for further prioritization. The DIABLO model achieved centroid distance error rates of 0.0625 (component 1) and 0.000 (component 2) for the metabolome data, and 0.1250 (component 1) and 0.1875 (component 2) for the microbiome data, indicating strong discriminatory performance across both blocks. This approach revealed strong associations between the relative abundances of Bacteroidales and Coriobacteriales and eight metabolites of interest, six of which possessed tryptophan-derived indolic structures, a class of metabolites known to mediate host-microbe and host-immune interactions.^57,58,59^ These included putative tryptophanol, tryptamine, 5-hydroxytryptophol, serotonin, and propionyl-histamine (Fig. 2D). This coordinated depletion of these metabolites in AMOX/CLAV-exposed pups suggests disruption of a microbial metabolic axis with known immunomodulatory roles.

Univariate testing showed significant reductions in (peak area) abundance of 5-hydroxy tryptophol, serotonin, and tryptophanol in AMOX/CLAV-exposed pups relative to mock-treated, PCV20-immunized controls (Wilcoxon, FDR-corrected *P* < 0.05) (Fig. 3A), mirroring the attenuated IgG vaccine responses observed in the same cohort (Fig. 1B). Propionyl-histamine levels were reduced in both antibiotic-exposed groups, with a greater reduction observed in the AMOX/CLAV cohort (Fig. 3A). Importantly, this indole-depletion phenotype was not dependent on vaccination, as subsetting to PBS-immunized pups showed similar reductions in the indolic-derived metabolites following antibiotic exposure (Fig. S4A). Thus, the metabolite shifts represent antibiotic-induced microbiome perturbation, rather than vaccine-driven metabolic changes.

**Figure 3.**
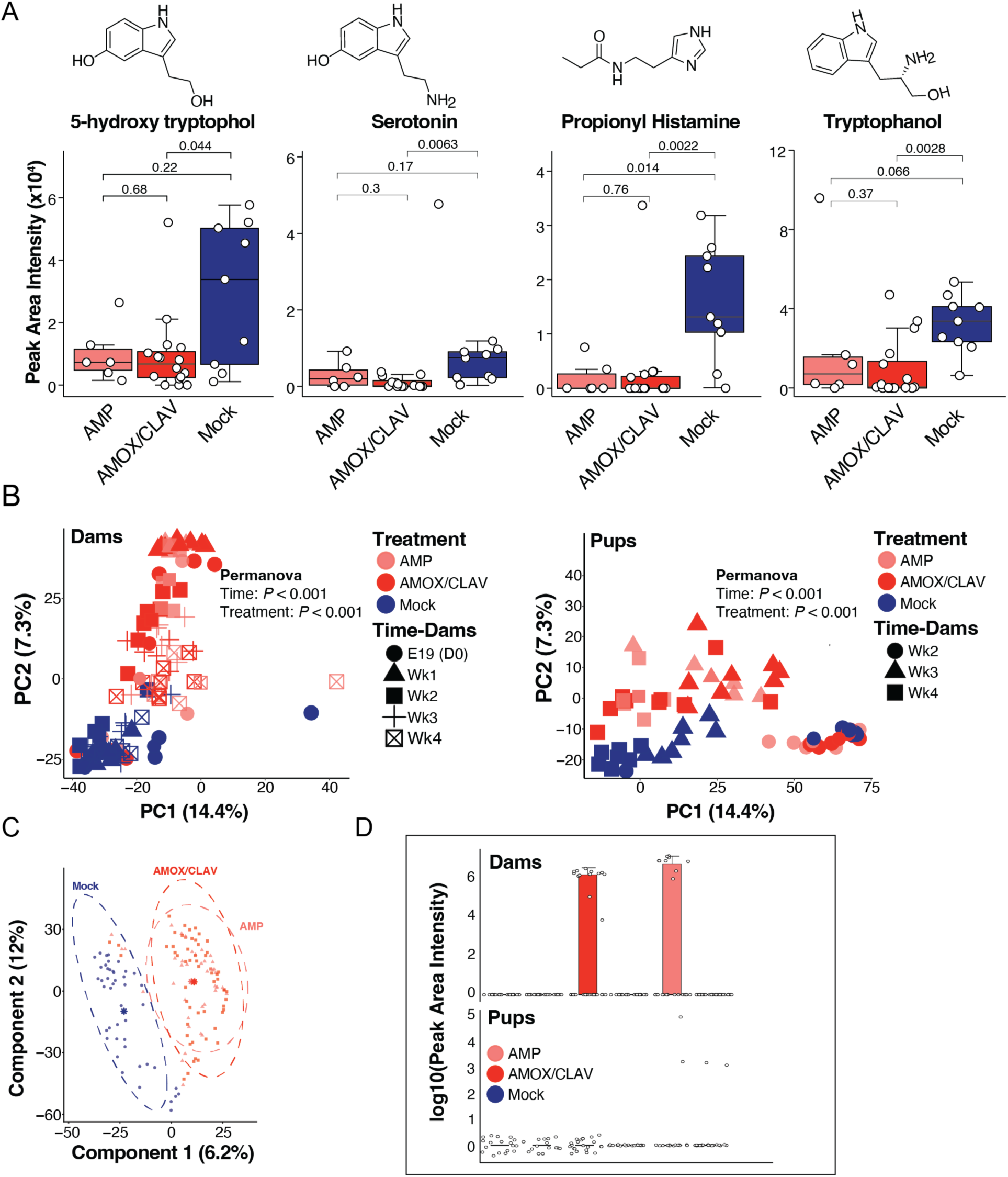
Metabolomics analysis of mice stratified by treatment condition. **(A)** Univariate analysis of selected molecular features 5-hydroxy tryptophol (feature ID 808), serotonin (feature ID805), propionyl histamine (feature ID 481), and tryptophanol (feature ID 2566). Levels of 5-hydroxy tryptophanol, serotonin, tryptophanol and propionyl histamine in pre-and post-immunized PCV20 Wk2-Wk4 pups that were indirectly exposed to antibiotics. Statistical significance was assessed using the Wilcoxon rank-sum test with benjamin-hochberg false discovery rate correction. The boxplots represent first (lower), interquartile range (IQR), and third (upper) quartile. **(B)** PCA of rclr-transformed metabolomic features revealed distinct clustering of mice based on antibiotic exposure. Separation was assessed using PERMANOVA, showing a significant effect of both treatment condition and time in pups and dams that were indirectly or directly exposed to antibiotics, respectively (PERMANOVA: treatment *P* < 0.001, time *P* < 0.001). **(C)** Partial Least Squares Discriminant Analysis (PLS-DA) of RCLR-transformed metabolomics features stratified by antibiotic treatment, with 95% confidence ellipses. Model performance evaluated by cross-validation yielded a balanced error rate of (BER) ≈ 0.43, indicating moderate classification accuracy. **(D)** Univariate analysis of detected features annotated as ampicillin or AMOX/CLAV presented as boxplots of summed log_10_(Peak area+1) transformed intensities in all treatment groups congregated at all time points. Ampicillin and AMOX/CLAV were detected in dams for their respective treatment groups but not detected in the pups’ feces.

### Global metabolomic signatures distinguish antibiotic-exposed and control groups

Unsupervised PCA revealed a clear separation between antibiotic-treated and mock-treated dams and offsprings, driven by both treatment and sampling time (PERMANOVA, dams, treatment: *P* = 0.001, R^2^ = 0.46233, F = 56.3932, time: *P* = 0.001, R^2^ = 0.12990, F = 7.9223) (PERMANOVA, pups, treatment: *P* = 0.001, R^2^ = 0.11880, F = 9.2615, time: *P* = 0.001 R^2^ = 0.55423, F = 43.2055; Fig. 3B). To interrogate treatment-specific metabolic signatures, we next applied a supervised PLS-DA model across all samples, comparing AMP-, AMOX/CLAV-, and mock-treated groups. This analysis again distinguished both antibiotic-treated cohorts from mock controls (Fig. 3C). Key discriminating metabolites were identified using variable importance in projection (VIP) scores from component 1, retaining annotated metabolites with VIP > 1 (Dataset 3). The top 50 VIP-ranked metabolites were visualized in a VIP dot plot and corresponding abundance heatmap, revealing that both antibiotic groups exhibited similar overall intensity patterns that were distinct from mock (Fig. 4). These features included putative bile acids, *N*-acyl lipids, dipeptides, and an acyl carnitine, among other small molecules. Notably, although AMP and AMOX/CLAV were detectable in dams during the treatment window (Fig. 3D, upper panel), neither antibiotic was detected in pup feces (Fig. 3D, lower panel) or serum (Dataset 4).

**Fig. 4.**
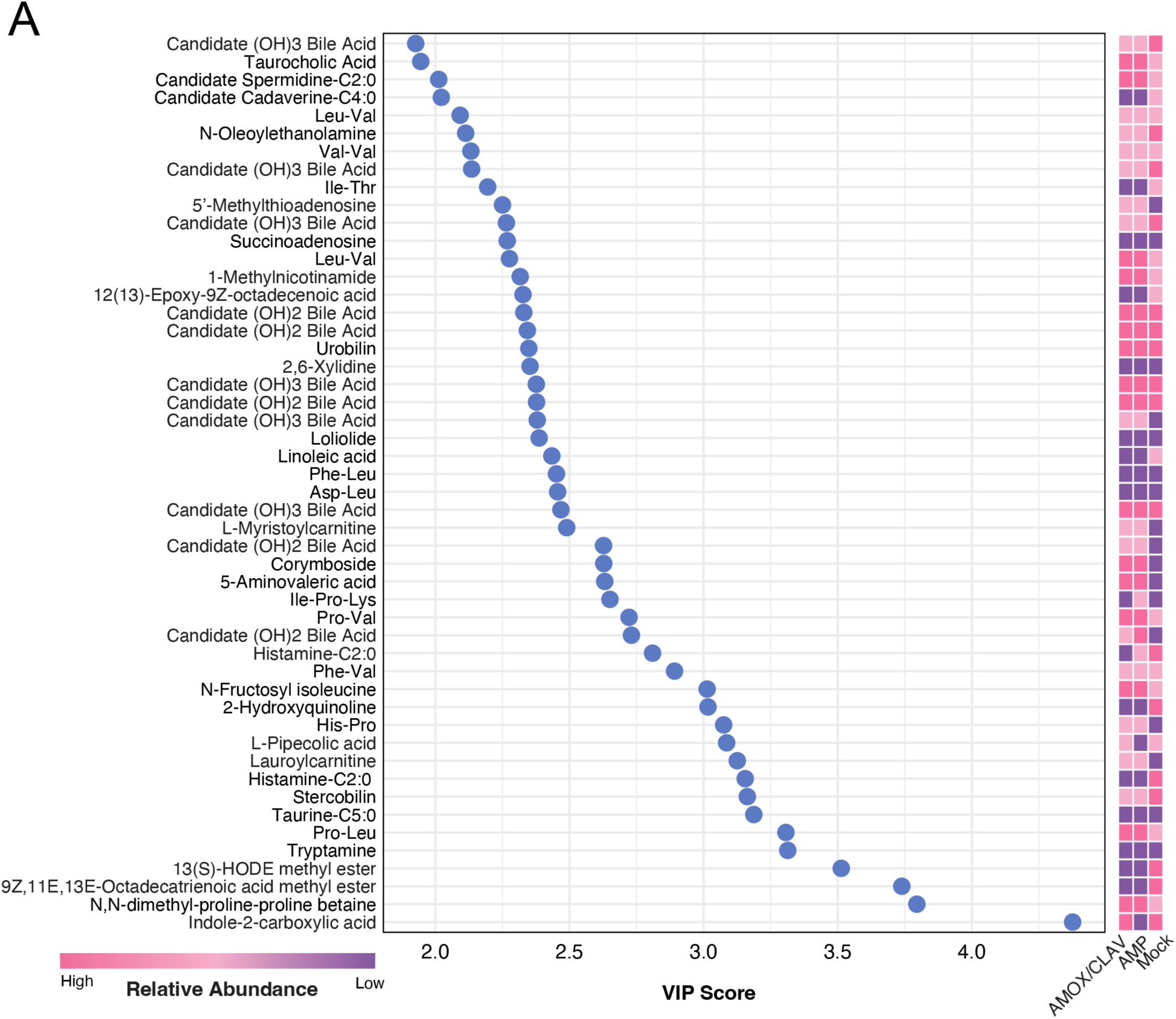
Heatmap and dot plot associated with metabolomics analysis of mice stratified by treatment condition. The plot and heatmap highlighting the top 50 annotated metabolites ranked by VIP scores. Heatmap represents the feature’s relative abundance detected across treatment groups. The annotations are based on MS/MS matching and represent Level 2 or 3 annotations according to the MSI.^88^ This means they have not been confirmed using standards and when a specific name is provided it is important to understand that other structural isomers are possible.

### Perturbation of microbial bile acid and *N*-acyl lipid metabolism in dams and offspring

Bile acid metabolism was examined by categorizing annotated features into three classes: non-conjugated bile acids, taurine-conjugated bile acids, and bile acids conjugated to alternative amines (e.g., leucine, phenylalanine, cysteamine). Antibiotic treatment elicited distinct and time-dependent shifts in bile acid composition in both dams and pups (Figs. S5A-B). At E19 and Wk1, both antibiotic-treated dam groups showed reduced levels of non-conjugated (free bile acids) and amine-conjugated bile acids (not including taurine and glycine) relative to mock controls (Figs. S5A-B). In AMOX/CLAV-treated dams, the abundance of non-conjugated bile acids increased significantly from E19/Wk1 to Wk2-4 (*P* < 0.0001), accompanied by a significant rise in amine-conjugated bile acids (*P* < 0.01) (Fig. S5A). AMP-treated dams showed a similar directional trend, though the changes did not reach statistical significance (Fig. S5A). In contrast, taurine-conjugated bile acids declined over time in both AMP (*P* < 0.01) and AMOX/CLAV (*P* < 0.001) groups (Fig. S5A); whereas mock-treated dams exhibited stable bile acid profiles across time (Fig. S5A). By Wk4, three weeks after cessation of antibiotic treatment, bile acid compositions converged across all dam groups (Fig. S5B). In antibiotic-treated pups, Wk2 samples showed a pronounced reduction in all bile acid classes relative to controls, with gradual recovery over time, trending toward mock-like levels Wk2 (Fig. S5B).

Beyond bile acid metabolism, antibiotic exposure also affected broader lipid pathways. Acylcarnitine levels were elevated in both dams and pups exposed to antibiotics (Fig. S6A). In contrast, microbially derived *N*-acyl lipids were reduced during antibiotic exposure, with partial recovery by Wk2 (Fig. S7A). Together, these data indicate that peripartum antibiotic exposure disrupts microbiota-controlled bile acid and lipid signaling pathways that support early immune maturation, providing a mechanistic link between AMOX/CLAV exposure and impaired vaccine responsiveness.

## DISCUSSION

Maternal antibiotic exposure during late gestation and lactation, particularly to broad-or extended-spectrum agents, perturbs early immune maturation and delays the establishment of effective vaccine-induced protection. Early-life immunity is tightly regulated by host-microbiota interactions, especially during the perinatal period when initial microbial colonization shapes developing immune networks and vaccine responsiveness.^60^ Using a murine model, we examined the consequences of maternal antibiotic exposure on neonatal responses to PCV20. We found that maternal treatment with AMOX/CLAV, but not AMP, impaired PCV20-induced antibody responses and neutrophil-mediated OPK activity in off-spring. Although these effects were transient (<8 weeks), the disruption occurred during a critical window of neonatal susceptibility, when delayed or suboptimal immune protection may increase infection risk.^61^ These findings are consistent with previous human^12^ and murine^10^ studies demonstrating that maternal antibiotic exposure can attenuate infant vaccine responses, particularly following broad-spectrum or high-dose antibiotic regimens.

The mechanistic basis for the loss of vaccine protection is likely multifactorial. Disruption of the maternal gut microbiota alters the pool of microorganisms available for vertical transmission at birth and during lactation, impairing neonatal gut colonization and the microbial signaling required for immune priming.^5,6,7,8,9^ Additionally, antibiotic-induced modulation of placental or mammary immunological signaling could contribute to altered fetal and neonatal immune programming.^15,62,63^

We further explored how maternal antibiotic exposure influences immune development through host-microbe pathways. Prior work has shown that perinatal antibiotics alter both maternal microbiome composition and metabolite profiles, with downstream effects on infant health.^64^ Consistent with these observations, maternal antibiotic treatment in our study led to coordinated shifts in the maternal gut microbiota and metabolome, with corresponding alterations observed in the offspring, supporting maternal-to-infant transmission of microbial community states. 16S rRNA sequencing revealed selective Bacteroidales and Coriobacteriales in AMOX/CLAV-exposed pups—taxa previously implicated in regulating inflammatory tone,^65^ T cell activation,^66^ and adaptive immune responses.^67^ Notably, this taxonomic disruption coincided with reduced PCV20-specific antibody responses, suggesting loss of these lineages may impair immune priming.

Both AMP and AMOX/CLAV exposure produced overlapping metabolic alterations, including elevated host-derived taurocholic acid^68^ and acylated carnitines,^69^ alongside depletion of microbiota-derived *N*-acyl lipids, which are known to support T cell differentiation and mucosal immune function^45^. These results reflect previous studies which found elevated levels of the same carnitines, including butyryl carnitine, decanoyl carnitine, dodecenoyl carnitine, and host-derived bile acids in perinatal exposure to AMP^64^. However, AMOX/CLAV induced additional, distinct changes characterized by depletion of non-conjugated bile acids,^68^ amine-conjugated bile acids, and tryptophan-derived indole metabolites. These metabolite classes signal through bile acid receptors and the aryl hydrocarbon receptor (AhR) to maintain epithelial integrity, regulate immune tone, and calibrate mucosal adaptive responses.^70,71,72^ Thus, AMOX/CLAV exposure disrupts a microbiota-dependent metabolic signaling network involving bile acids, *N*-acyl lipids, and indole-AhR ligands—pathways that collectively contribute to effective vaccine-induced immunity.

For example, taurocholic acid, which was elevated in both antibiotic treatment groups, has been shown to promote the expansion of pathogenic bacteria *in vivo*.^73^ Antibiotic exposure also resulted in depletion of bile amidates, microbial products implicated in gut-immune communication,^74,75^ and accompanied reduced levels of microbially-derived (non-conjugated) bile acids and elevated levels of host-conjugated bile acids.^76^ In parallel, *N*-acyl lipids, which were depleted in both AMP-and AMP/CLAV-treated groups, are produced by >70 commensal species and contribute to T cell differentiation and mucosal immune function.^45^ Although short-chain fatty acids (SCFAs) were not directly quantified in our data set, the loss of *N*-acyl lipids is consistent with reduced SCFA biosynthetic capacity, a pathway with well-established roles in regulating immune homeostasis and vaccine responsiveness^77^.

Integrative multi-omics analysis revealed strong correlations between immune-associated microbial taxa and tryptophan-derived indole metabolites. Several such metabolites, including indolelactic acid (ILA), indolepropionic acid (IPA), and indoleacetic acid (IAA), are produced by commensal bacteria in the infant gut.^78^ Signaling through AhR, these indole derivatives can influence mucosal barrier integrity, T follicular helper (Tfh) cell differentiation, germinal center formation, and cytokine signaling.^79,80,81^ Additional tryptophan-derived metabolites detected here—including serotonin, tryptamine, tryptophol, and 5-hydroxy tryptophol, as well as two isobaric tryptophanol species—were significantly depleted in AMOX/CLAV-exposed infants. *Bifidobacterium* and *Bacteroides* spp. are recognized producers of these metabolites.^82^ Taken together, these findings suggest that disruption of Bacteroidales-mediated tryptophan metabolism decreases the availability of AhR-active microbial metabolites, leading to impaired mucosal immune signaling and suboptimal germinal center responses, and ultimately reduced vaccine-specific antibody function.

Unnecessary exposure to broad-spectrum antibiotics remains widespread^83^, and AMOX/CLAV is currently being evaluated as an orally administered prophylactic agent for reducing GBS and other infection-related risks^84,85,86,87^. We demonstrate that AMOX/CLAV selectively perturbs this microbial–metabolic axis, whereas AMP does not, thereby providing a mechanistic basis for sub-optimal neonatal vaccine responsiveness observed *in vivo*. These findings suggest that minimizing unnecessary exposure to AMOX/CLAV use during pregnancy and lactation will help to preserve early-life vaccine responsiveness and immune function. Future studies will be needed to identify the strain-level contributors within Bacteroidales and Coriobacteriales, determine the immune cell populations most affected (e.g., Tfh, Treg, and germinal center B cells), and evaluate whether targeted microbial or metabolite supplementation can restore vaccine responsiveness during the neonatal immune development window.

## LIMITATIONS

This study examined the metabolic mechanisms underlying microbially-mediated immunity in early life, leveraging high-throughput sequencing and high-resolution mass spectrometry for metabolite identification. Several limitations merit consideration. Metabolites were annotated by MS/MS spectral matching (metabolomics standard initiative level 2)^88^, against GNPS spectral libraries. Although AMP and AMOX/CLAV were detected in the feces of antibiotic treated dams, they were not detected in pup feces or serum. This absence may reflect limited transfer of antibiotics to offspring and/or analytical sensitivity constraints. Future studies should incorporate higher-sensitivity instrumentation and targeted quantification approaches to determine whether these antibiotics are transmitted through breast milk. While oral AMP or AMOX/CLAV dosing effectively perturbed the maternal gut microbiota, it does not fully mirror the intravenous regimens commonly used in clinical intrapartum antibiotic prophylaxis; consequently, systemic exposure and transfer to offsprings may differ. Nevertheless, this model was suitable for interrogating microbiome-mediated effects on early immune development while maintaining offspring viability, as minimizing maternal stress reduces the risk of cannibalism and pup loss, an important consideration in neonatal studies. Finally, although the sample size was limited, the pronounced effects observed in this study replicate or align with multiple prior studies of antibiotic-induced gut microbiome perturbations, thereby strengthening confidence in our conclusions.

## SUPPLEMENTAL INFORMATION

The Supplemental information contains Supplementary figures (**Figures S1–S7**), and description of additional supplementary files (dataset legends). Expanded metabolomics and microbiome data associated with this manuscript are included in Datasets MSV000097137.

## RESOURCES AND DATA AVAILABILITY

All data required to evaluate the paper’s conclusions are included in the manuscript and/or Supporting Information. Metabolomics and microbiome data have been deposited in GNPS/MassIVE as well as Qiita with a dataset identifier MSV000097137 and 15920, respectively. The EBI accession number is ERP186116. Source code is available under the assigned DOIs <u>10.5281/zenodo.17726094</u> and <u>10.5281/zenodo.17728254</u>.

## ACKNOWLEDGMENT

This work was supported by the Eunice Kennedy Shriver National Institute of Child Health Development Maternal Pediatric Precision in Therapeutics (MPRINT) grant (P50HD106463), National Institute of General Medical Sciences (R01GM107550), and the National Science Foundation (DBI-2152526). We are grateful for services provided by the University of California, San Diego, Center for Microbiome Innovation.

## AUTHOR CONTRIBUTIONS

F.A., E.S., V.N., G.L. conceived and designed the study and the experiments. E.S., F.A., A.K. C.-M.T., H.N., and D.S. conducted animal experiments; E.S. F.A., V.D., J.Z. analyzed data; S.Z. and K.E.K contributed code for processing the metabolomics and microbiome data, which was analyzed by J.Z and V.D; S.M.T and P.C.D contributed with microbiome and metabolome analytic tools and intellectual input; F. A., E.S., V.D., J.Z., wrote the manuscript, which all co-authors reviewed and approved.

## REFERENCES

1. Fairlie T, Zell ER, Schrag S. Effectiveness of intrapartum antibiotic prophylaxis for prevention of early-onset group B streptococcal disease. Obstet Gynecol 2013; 121:570–7.

2. Boyer KM, Gotoff SP. Prevention of early-onset neonatal group B streptococcal disease with selective intrapartum chemoprophylaxis. N Engl J Med 1986; 314:1665–9.

3. Azad MB, Konya T, Persaud RR, Guttman DS, Chari RS, Field CJ, Sears MR, Mandhane PJ, Turvey SE, Subbarao P, et al. Impact of maternal intrapartum antibiotics, method of birth and breastfeeding on gut microbiota during the first year of life: a prospective cohort study. BJOG 2016; 123:983–93.

4. Persaud RR, Azad MB, Chari RS, Sears MR, Becker AB, Kozyrskyj AL, CHILD Study Investigators. Perinatal antibiotic exposure of neonates in Canada and associated risk factors: a population-based study. J Matern Fetal Neonatal Med 2015; 28:1190–5.

5. Ignacio A, Czyz S, McCoy KD. Early life microbiome influences on development of the mucosal innate immune system. Semin Immunol 2024; 73:101885.

6. Jain N. The early life education of the immune system: Moms, microbes and (missed) opportunities. Gut Microbes 2020; 12:1824564.

7. Paucar Iza YA, Brown CC. Early life imprinting of intestinal immune tolerance and tissue homeostasis. Immunol Rev 2024; 323:303–15.

8. Al Nabhani Z, Eberl G. Imprinting of the immune system by the microbiota early in life. Mucosal Immunol 2020; 13:183–9.

9. Vangay P, Ward T, Gerber JS, Knights D. Antibiotics, pediatric dysbiosis, and disease. Cell Host Microbe 2015; 17:553–64.

10. Lynn MA, Tumes DJ, Choo JM, Sribnaia A, Blake SJ, Leong LEX, Young GP, Marshall HS, Wesselingh SL, Rogers GB, et al. Early-Life Antibiotic-Driven Dysbiosis Leads to Dysregulated Vaccine Immune Responses in Mice. Cell Host Microbe 2018; 23:653–60.e5.

11. Ryan FJ, Clarke M, Lynn MA, Benson SC, McAlister S, Giles LC, Choo JM, Rossouw C, Ng YY, Semchenko EA, et al. Bifidobacteria support optimal infant vaccine responses. Nature 2025; 641:456–64.

12. Chapman TJ, Pham M, Bajorski P, Pichichero ME. Antibiotic Use and Vaccine Antibody Levels. Pediatrics [Internet] 2022; 149. Available from: 10.1542/peds.2021-052061

13. Shaffer M, Best K, Tang C, Liang X, Schulz S, Gonzalez E, White CH, Wyche TP, Kang J, Wesseling H, et al. Very early life microbiome and metabolome correlates with primary vaccination variability in children. mSystems 2023; 8:e0066123.

14. Pichichero ME, Xu L, Gonzalez E, Pham M, Kaur R. Variability of Vaccine Responsiveness in Young Children. J Infect Dis 2024; 229:1856–65.

15. Pietrasanta C, Carlosama C, Lizier M, Fornasa G, Jost TR, Carloni S, Giugliano S, Silvestri A, Brescia P, De Ponte Conti B, et al. Prenatal antibiotics reduce breast milk IgA and induce dysbiosis in mouse offspring, increasing neonatal susceptibility to bacterial sepsis. Cell Host Microbe 2024; 32:2178–94.e6.

16. Thomas SP, Denizer E, Zuffa S, Best BM, Bode L, Chambers CD, Dorrestein PC, Liu GY, Momper JD, Nizet V, et al. Transfer of antibiotics and their metabolites in human milk: Implications for infant health and microbiota. Pharmacotherapy 2023; 43:442–51.

17. Orwa SA, Gudnadottir U, Boven A, Pauwels I, Versporten A, Vlieghe E, Brusselaers N. Global prevalence of antibiotic consumption during pregnancy: A systematic review and meta-analysis. J Infect 2024; 89:106189.

18. Lebeaux RM, Karalis DB, Lee J, Whitehouse HC, Madan JC, Karagas MR, Hoen AG. The association between early life antibiotic exposure and the gut resistome of young children: a systematic review. Gut Microbes 2022; 14:2120743.

19. Räty S, Ollila H, Turta O, Pärtty A, Peltola V, Lagström H, Lempainen J, Rautava S. Neonatal and early infancy antibiotic exposure is associated with childhood atopic dermatitis, wheeze and asthma. Eur J Pediatr 2024; 183:5191–202.

20. Adami AJ, Bracken SJ, Guernsey LA, Rafti E, Maas KR, Graf J, Matson AP, Thrall RS, Schramm CM. Early-life antibiotics attenuate regulatory T cell generation and increase the severity of murine house dust mite-induced asthma. Pediatr Res 2018; 84:426–34.

21. Livanos AE, Greiner TU, Vangay P, Pathmasiri W, Stewart D, McRitchie S, Li H, Chung J, Sohn J, Kim S, et al. Antibiotic-mediated gut microbiome perturbation accelerates development of type 1 diabetes in mice. Nat Microbiol 2016; 1:16140.

22. Vijay-Kumar M, Aitken JD, Carvalho FA, Cullender TC, Mwangi S, Srinivasan S, Sitaraman SV, Knight R, Ley RE, Gewirtz AT. Metabolic syndrome and altered gut microbiota in mice lacking Toll-like receptor 5. Science 2010; 328:228–31.

23. Turnbaugh PJ, Ley RE, Mahowald MA, Magrini V, Mardis ER, Gordon JI. An obesity-associated gut microbiome with increased capacity for energy harvest. Nature 2006; 444:1027–31.

24. Russell SL, Gold MJ, Hartmann M, Willing BP, Thorson L, Wlodarska M, Gill N, Blanchet M-R, Mohn WW, McNagny KM, et al. Early life antibiotic-driven changes in microbiota enhance susceptibility to allergic asthma. EMBO Rep 2012; 13:440–7.

25. Jensen ET, Svane HM, Erichsen R, Kurt G, Heide-Jorgensen U, Sorensen HT, Dellon ES. Maternal and Infant Antibiotic and Acid Suppressant Use and Risk of Eosinophilic Esophagitis. JAMA Pediatr 2023; 177:1285–93.

26. Jawad AB, Jansson S, Wewer V, Malham M. Early life oral antibiotics are associated with pediatric-onset inflammatory bowel disease-A nationwide study. J Pediatr Gastroenterol Nutr 2023; 77:366–72.

27. Thacker N, Duncanson K, Eslick GD, Dutt S, O’Loughlin EV, Hoedt EC, Collins CE. Antibiotics, passive smoking, high socioeconomic status and sweetened foods contribute to the risk of paediatric inflammatory bowel disease: A systematic review with meta-analysis. J Pediatr Gastroenterol Nutr 2024; 79:610–21.

28. Räisänen LK, Kääriäinen SE, Sund R, Engberg E, Viljakainen HT, Kolho K-L. Antibiotic exposures and the development of pediatric autoimmune diseases: a register-based case-control study. Pediatr Res 2023; 93:1096–104.

29. Benitez AJ, Tanes C, Friedman ES, Zackular JP, Ford E, Gerber JS, DeRusso PA, Kelly A, Li H, Elovitz MA, et al. Antibiotic exposure is associated with minimal gut microbiome perturbations in healthy term infants. Microbiome 2025; 13:21.

30. Schoch JJ, Satcher KG, Garvan CW, Monir RL, Neu J, Lemas DJ. Association between early life antibiotic exposure and development of early childhood atopic dermatitis. JAAD Int 2023; 10:68–74.

31. Jin S, Zhao D, Cai C, Song D, Shen J, Xu A, Qiao Y, Ran Z, Zheng Q. Low-dose penicillin exposure in early life decreases Th17 and the susceptibility to DSS colitis in mice through gut microbiota modification. Sci Rep 2017; 7:43662.

32. Zwittink RD, van Zoeren-Grobben D, Renes IB, van Lingen RA, Norbruis OF, Martin R, Groot Jebbink LJ, Knol J, Belzer C. Dynamics of the bacterial gut microbiota in preterm and term infants after intravenous amoxicillin/ceftazidime treatment. BMC Pediatr 2020; 20:195.

33. Korpela K, Salonen A, Saxen H, Nikkonen A, Peltola V, Jaakkola T, de Vos W, Kolho K-L. Antibiotics in early life associate with specific gut microbiota signatures in a prospective longitudinal infant cohort. Pediatr Res 2020; 88:438–43.

34. Van Daele E, Kamphorst K, Vlieger AM, Hermes G, Milani C, Ventura M, Belzer C, Smidt H, van Elburg RM, Knol J. Effect of antibiotics in the first week of life on faecal microbiota development. Arch Dis Child Fetal Neonatal Ed 2022; 107:603–10.

35. Barnett DJM, Endika MF, Klostermann CE, Gu F, Thijs C, Nauta A, Schols HA, Smidt H, Arts ICW, Penders J. Human milk oligosaccharides, antimicrobial drugs, and the gut microbiota of term neonates: observations from the KOALA birth cohort study. Gut Microbes 2023; 15:2164152.

36. Reyman M, van Houten MA, Watson RL, Chu MLJN, Arp K, de Waal WJ, Schiering I, Plötz FB, Willems RJL, van Schaik W, et al. Effects of early-life antibiotics on the developing infant gut microbiome and resistome: a randomized trial. Nat Commun 2022; 13:893.

37. Mangin I, Suau A, Gotteland M, Brunser O, Pochart P. Amoxicillin treatment modifies the composition of Bifidobacterium species in infant intestinal microbiota. Anaerobe 2010; 16:433–8.

38. Huda MN, Ahmad SM, Alam MJ, Khanam A, Kalanetra KM, Taft DH, Raqib R, Underwood MA, Mills DA, Stephensen CB. Abundance in Early Infancy and Vaccine Response at 2 Years of Age. Pediatrics [Internet] 2019; 143. Available from: 10.1542/peds.2018-1489

39. Caldera JR, Tsai C-M, Trieu D, Gonzalez C, Hajam IA, Du X, Lin B, Liu GY. The characteristics of pre-existing humoral imprint determine efficacy of S. aureus vaccines and support alternative vaccine approaches. Cell Rep Med 2024; 5:101360.

40. Chambers MC, Maclean B, Burke R, Amodei D, Ruderman DL, Neumann S, Gatto L, Fischer B, Pratt B, Egertson J, et al. A cross-platform toolkit for mass spectrometry and proteomics. Nat Biotechnol 2012; 30:918–20.

41. Schmid R, Heuckeroth S, Korf A, Smirnov A, Myers O, Dyrlund TS, Bushuiev R, Murray KJ, Hoffmann N, Lu M, et al. Integrative analysis of multimodal mass spectrometry data in MZmine 3. Nat Biotechnol 2023; 41:447–9.

42. Nothias L-F, Petras D, Schmid R, Dührkop K, Rainer J, Sarvepalli A, Protsyuk I, Ernst M, Tsugawa H, Fleischauer M, et al. Feature-based molecular networking in the GNPS analysis environment. Nat Methods 2020; 17:905–8.

43. Mohanty I, Mannochio-Russo H, Schweer JV, El Abiead Y, Bittremieux W, Xing S, Schmid R, Zuffa S, Vasquez F, Muti VB, et al. The underappreciated diversity of bile acid modifications. Cell 2024; 187:1801–18.e20.

44. Mohanty I, Xing S, Castillo V, Agongo J, Patan A, El Abiead Y, Mannochio-Russo H, Zuffa S, Zemlin J, Tronel A, et al. MS/MS mass spectrometry filtering tree for bile acid isomer annotation [Internet]. bioRxiv 2025; Available from: http://biorxiv.org/lookup/doi/10.1101/2025.03.04.641505

45. Mannochio-Russo H, Charron-Lamoureux V, van Faassen M, Lamichhane S, Gonçalves Nunes WD, Deleray V, Ayala AV, Tanaka Y, Patan A, Vittali K, et al. The microbiome diversifies long-to short-chain fatty acid-derived N-acyl lipids. Cell 2025; 188:4154–69.e19.

46. Marotz C, Belda-Ferre P, Ali F, Das P, Huang S, Cantrell K, Jiang L, Martino C, Diner RE, Rahman G, et al. SARS-CoV-2 detection status associates with bacterial community composition in patients and the hospital environment. Microbiome 2021; 9:132.

47. Das S, Dash HR. Microbial Diversity in the Genomic Era. Academic Press; 2018.

48. Walters W, Hyde ER, Berg-Lyons D, Ackermann G, Humphrey G, Parada A, Gilbert JA, Jansson JK, Caporaso JG, Fuhrman JA, et al. Improved Bacterial 16S rRNA Gene (V4 and V4-5) and Fungal Internal Transcribed Spacer Marker Gene Primers for Microbial Community Surveys. mSystems [Internet] 2016; 1. Available from: 10.1128/mSystems.00009-15

49. Minich JJ, Humphrey G, Benitez RAS, Sanders J, Swafford A, Allen EE, Knight R. High-Throughput Miniaturized 16S rRNA Amplicon Library Preparation Reduces Costs while Preserving Microbiome Integrity. mSystems [Internet] 2018; 3. Available from: 10.1128/mSystems.00166-18

50. Marotz C, Sharma A, Humphrey G, Gottel N, Daum C, Gilbert JA, Eloe-Fadrosh E, Knight R. Triplicate PCR reactions for 16S rRNA gene amplicon sequencing are unnecessary. Biotechniques 2019; 67:29–32.

51. Gonzalez A, Navas-Molina JA, Kosciolek T, McDonald D, Vázquez-Baeza Y, Ackermann G, DeReus J, Janssen S, Swafford AD, Orchanian SB, et al. Qiita: rapid, web-enabled microbiome meta-analysis. Nat Methods 2018; 15:796–8.

52. Amir A, McDonald D, Navas-Molina JA, Kopylova E, Morton JT, Zech Xu Z, Kightley EP, Thompson LR, Hyde ER, Gonzalez A, et al. Deblur Rapidly Resolves Single-Nucleotide Community Sequence Patterns. mSystems [Internet] 2017; 2. Available from: 10.1128/mSystems.00191-16

53. Bolyen E, Rideout JR, Dillon MR, Bokulich NA, Abnet CC, Al-Ghalith GA, Alexander H, Alm EJ, Arumugam M, Asnicar F, et al. Reproducible, interactive, scalable and extensible microbiome data science using QIIME 2. Nat Biotechnol 2019; 37:852–7.

54. McDonald D, Jiang Y, Balaban M, Cantrell K, Zhu Q, Gonzalez A, Morton JT, Nicolaou G, Parks DH, Karst SM, et al. Greengenes2 unifies microbial data in a single reference tree. Nat Biotechnol 2024; 42:715–8.

55. Fernandes AD, Reid JN, Macklaim JM, McMurrough TA, Edgell DR, Gloor GB. Unifying the analysis of high-throughput sequencing datasets: characterizing RNA-seq, 16S rRNA gene sequencing and selective growth experiments by compositional data analysis. Microbiome 2014; 2:15.

56. Singh A, Shannon CP, Gautier B, Rohart F, Vacher M, Tebbutt SJ, Lê Cao K-A. DIABLO: an integrative approach for identifying key molecular drivers from multi-omics assays. Bioinformatics 2019; 35:3055–62.

57. Fiore A, Murray PJ. Tryptophan and indole metabolism in immune regulation. Curr Opin Immunol 2021; 70:7–14.

58. Gao J, Xu K, Liu H, Liu G, Bai M, Peng C, Li T, Yin Y. Impact of the Gut Microbiota on Intestinal Immunity Mediated by Tryptophan Metabolism. Front Cell Infect Microbiol 2018; 8:13.

59. Li S. Modulation of immunity by tryptophan microbial metabolites. Front Nutr 2023; 10:1209613.

60. Lynn DJ, Benson SC, Lynn MA, Pulendran B. Modulation of immune responses to vaccination by the microbiota: implications and potential mechanisms. Nat Rev Immunol 2022; 22:33–46.

61. Pieren DKJ, Boer MC, de Wit J. The adaptive immune system in early life: The shift makes it count. Front Immunol 2022; 13:1031924.

62. Gonzalez-Perez G, Hicks AL, Tekieli TM, Radens CM, Williams BL, Lamousé-Smith ESN. Maternal Antibiotic Treatment Impacts Development of the Neonatal Intestinal Microbiome and Antiviral Immunity. J Immunol 2016; 196:3768–79.

63. Benner M, Lopez-Rincon A, Thijssen S, Garssen J, Ferwerda G, Joosten I, van der Molen RG, Hogenkamp A. Antibiotic Intervention Affects Maternal Immunity During Gestation in Mice. Front Immunol 2021; 12:685742.

64. Zuffa S, Thomas SP, Mohanty I, El Abiead Y, Deleray V, Kvitne KE, Kousha A, Suzuki E, Tsai CM, Nguyen G, et al. Influence of perinatal ampicillin exposure on maternal fecal microbial and metabolic profiles [Internet]. bioRxiv 2025 [cited 2025 Oct 22];:2025.06.30.662372. Available from: https://pmc.ncbi.nlm.nih.gov/articles/PMC12236745/

65. Kuhn KA, Schulz HM, Regner EH, Severs EL, Hendrickson JD, Mehta G, Whitney AK, Ir D, Ohri N, Robertson CE, et al. Bacteroidales recruit IL-6-producing intraepithelial lymphocytes in the colon to promote barrier integrity. Mucosal Immunol 2018; 11:357–68.

66. Vétizou M, Pitt JM, Daillère R, Lepage P, Waldschmitt N, Flament C, Rusakiewicz S, Routy B, Roberti MP, Duong CPM, et al. Anticancer immunotherapy by CTLA-4 blockade relies on the gut microbiota. Science 2015; 350:1079–84.

67. Riazati N, Kable ME, Stephensen CB. Association of intestinal bacteria with immune activation in a cohort of healthy adults. Microbiol Spectr 2023; 11:e0102723.

68. Quinn RA, Melnik AV, Vrbanac A, Fu T, Patras KA, Christy MP, Bodai Z, Belda-Ferre P, Tripathi A, Chung LK, et al. Global chemical effects of the microbiome include new bile-acid conjugations. Nature 2020; 579:123–9.

69. McCann MR, George De la Rosa MV, Rosania GR, Stringer KA. L-Carnitine and Acylcarnitines: Mitochondrial Biomarkers for Precision Medicine. Metabolites 2021; 11:51.

70. Godlewska U, Bulanda E, Wypych TP. Bile acids in immunity: Bidirectional mediators between the host and the microbiota. Front Immunol 2022; 13:949033.

71. Masse KE, Lu VB. Short-chain fatty acids, secondary bile acids and indoles: gut microbial metabolites with effects on enteroendocrine cell function and their potential as therapies for metabolic disease. Front Endocrinol 2023; 14:1169624.

72. Yang W, Cong Y. Gut microbiota-derived metabolites in the regulation of host immune responses and immune-related inflammatory diseases. Cell Mol Immunol 2021; 18:866–77.

73. Devkota S, Wang Y, Musch MW, Leone V, Fehlner-Peach H, Nadimpalli A, Antonopoulos DA, Jabri B, Chang EB. Dietary-fat-induced taurocholic acid promotes pathobiont expansion and colitis in Il10-/-mice. Nature 2012; 487:104–8.

74. Guzior DV, Okros M, Shivel M, Armwald B, Bridges C, Fu Y, Martin C, Schilmiller AL, Miller WM, Ziegler KM, et al. Bile salt hydrolase acyltransferase activity expands bile acid diversity. Nature 2024; 626:852–8.

75. Mohanty I, Allaband C, Mannochio-Russo H, El Abiead Y, Hagey LR, Knight R, Dorrestein PC. The changing metabolic landscape of bile acids – keys to metabolism and immune regulation. Nature Reviews Gastroenterology & Hepatology 2024; 21:493–516.

76. Brestoff JR, Artis D. Commensal bacteria at the interface of host metabolism and the immune system. Nat Immunol 2013; 14:676–84.

77. Mann ER, Lam YK, Uhlig HH. Short-chain fatty acids: linking diet, the microbiome and immunity. Nat Rev Immunol 2024; 24:577–95.

78. Laursen MF, Sakanaka M, von Burg N, Mörbe U, Andersen D, Moll JM, Pekmez CT, Rivollier A, Michaelsen KF, Mølgaard C, et al. Bifidobacterium species associated with breastfeeding produce aromatic lactic acids in the infant gut. Nat Microbiol 2021; 6:1367–82.

79. Hezaveh K, Shinde RS, Klötgen A, Halaby MJ, Lamorte S, Ciudad MT, Quevedo R, Neufeld L, Liu ZQ, Jin R, et al. Tryptophan-derived microbial metabolites activate the aryl hydrocarbon receptor in tumor-associated macrophages to suppress anti-tumor immunity. Immunity 2022; 55:324–40.e8.

80. Gutiérrez-Vázquez C, Quintana FJ. Regulation of the Immune Response by the Aryl Hydrocarbon Receptor. Immunity 2018; 48:19–33.

81. Zelante T, Iannitti RG, Cunha C, De Luca A, Giovannini G, Pieraccini G, Zecchi R, D’Angelo C, Massi-Benedetti C, Fallarino F, et al. Tryptophan catabolites from microbiota engage aryl hydrocarbon receptor and balance mucosal reactivity via interleukin-22. Immunity 2013; 39:372–85.

82. Zhang N-N, Jiang Z-M, Li S-Z, Yang X, Liu E-H. Evolving interplay between natural products and gut microbiota. Eur J Pharmacol 2023; 949:175557.

83. Cardetti M, Rodríguez S, Sola A. Use (and abuse) of antibiotics in perinatal medicine. An Pediatr (Engl Ed) 2020; 93:207.e1–207.e7.

84. Kenyon SL, Taylor DJ, Tarnow-Mordi W, ORACLE Collaborative Group. Broad-spectrum antibiotics for preterm, prelabour rupture of fetal membranes: the ORACLE I randomised trial. ORACLE Collaborative Group. Lancet 2001; 357:979–88.

85. Yudin MH, van Schalkwyk J, Van Eyk N. No. 233-Antibiotic Therapy in Preterm Premature Rupture of the Membranes. J Obstet Gynaecol Can 2017; 39:e207–12.

86. Knight M, Chiocchia V, Partlett C, Rivero-Arias O, Hua X, Hinshaw K, Tuffnell D, Linsell L, Juszczak E, ANODE collaborative group. Prophylactic antibiotics in the prevention of infection after operative vaginal delivery (ANODE): a multicentre randomised controlled trial. Lancet 2019; 393:2395–403.

87. Keij FM, Kornelisse RF, Hartwig NG, van der Sluijs-Bens J, van Beek RHT, van Driel A, van Rooij LGM, van Dalen-Vink I, Driessen GJA, Kenter S, et al. Efficacy and safety of switching from intravenous to oral antibiotics (amoxicillin-clavulanic acid) versus a full course of intravenous antibiotics in neonates with probable bacterial infection (RAIN): a multicentre, randomised, open-label, non-inferiority trial. Lancet Child Adolesc Health 2022; 6:799–809.

88. Fiehn O, Robertson D, Griffin J, van der Werf M, Nikolau B, Morrison N, Sumner LW, Goodacre R, Hardy NW, Taylor C, et al. The metabolomics standards initiative (MSI). Metabolomics 2007; 3:175–8.

