## Supplementary figures and images for "Effect of Perinatal Ampicillin or Amoxicillin/Clavulanate Exposure on Maternal and Infant Gut Microbiome, Metabolome, and Infant Responses to the 20-valent Pneumococcal Conjugate Vaccine"

### Supplementary Figure 1

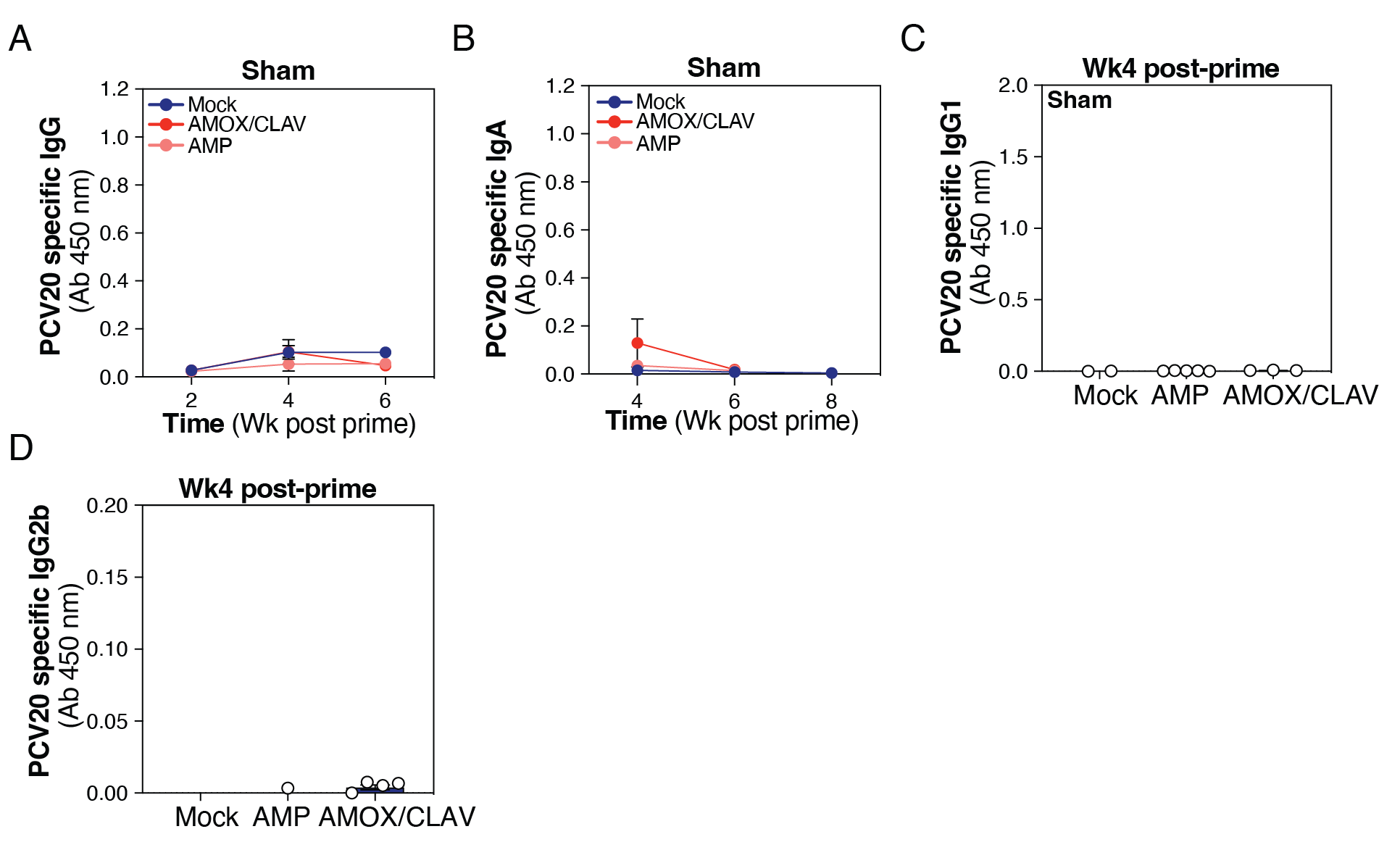

### Supplementary Figure 2

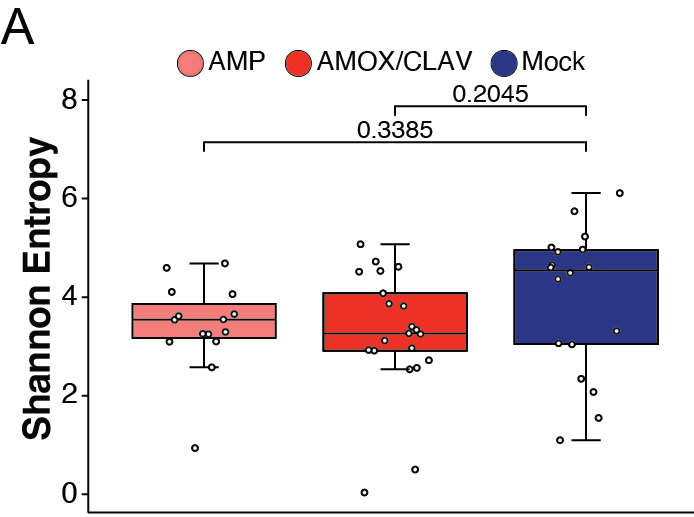

### Supplementary Figure 3

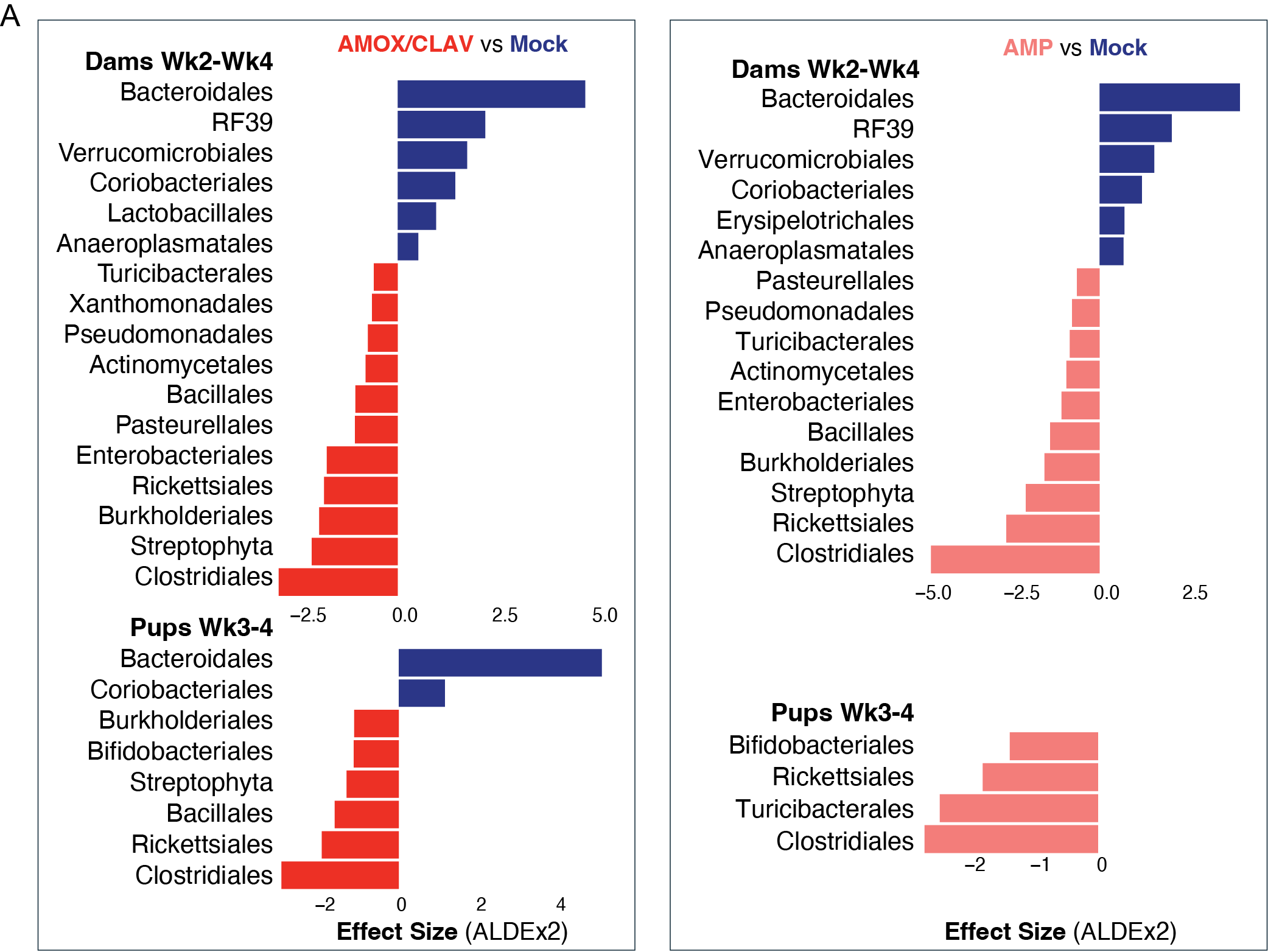

### Supplementary Figure 4

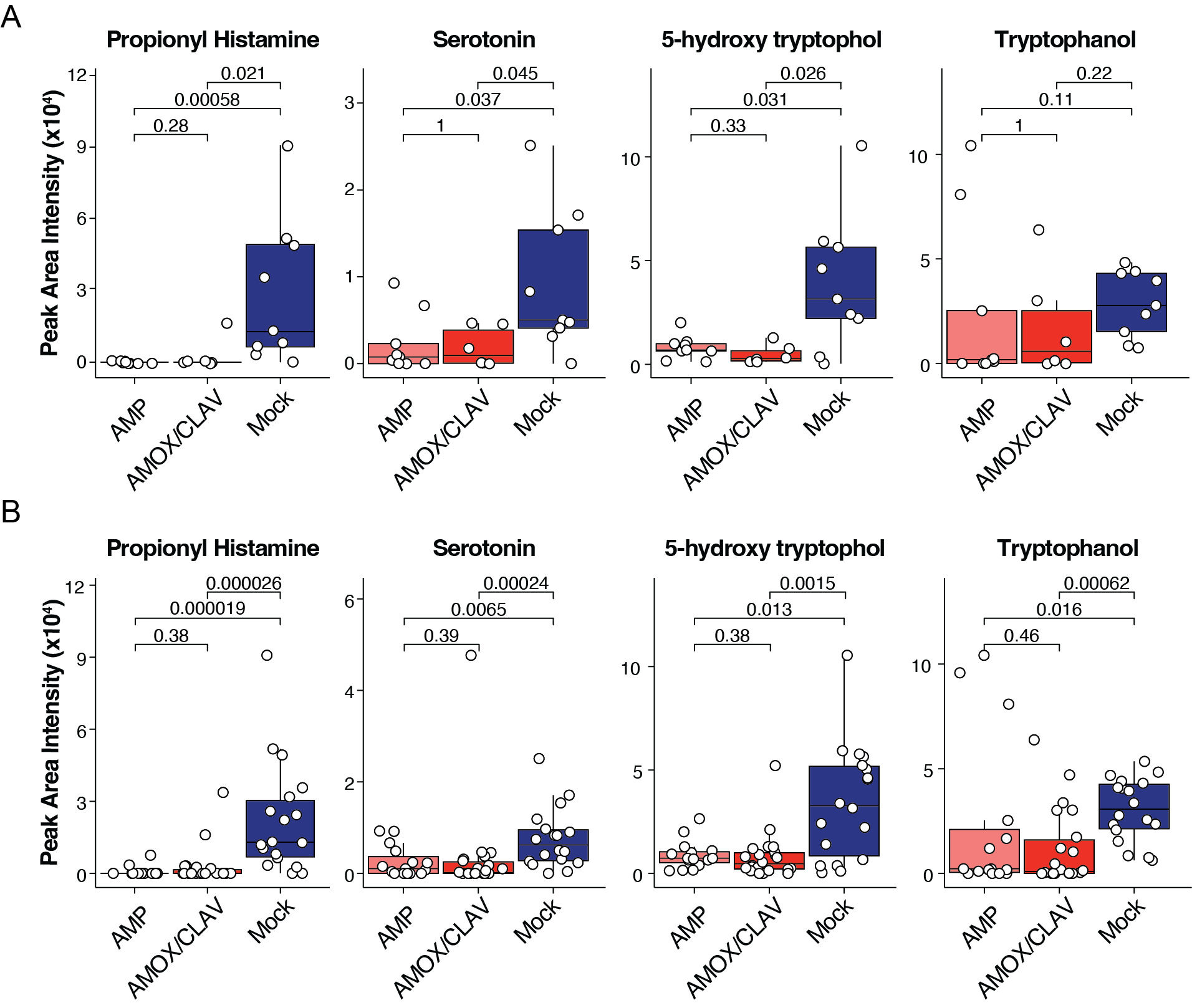

### Supplementary Figure 5

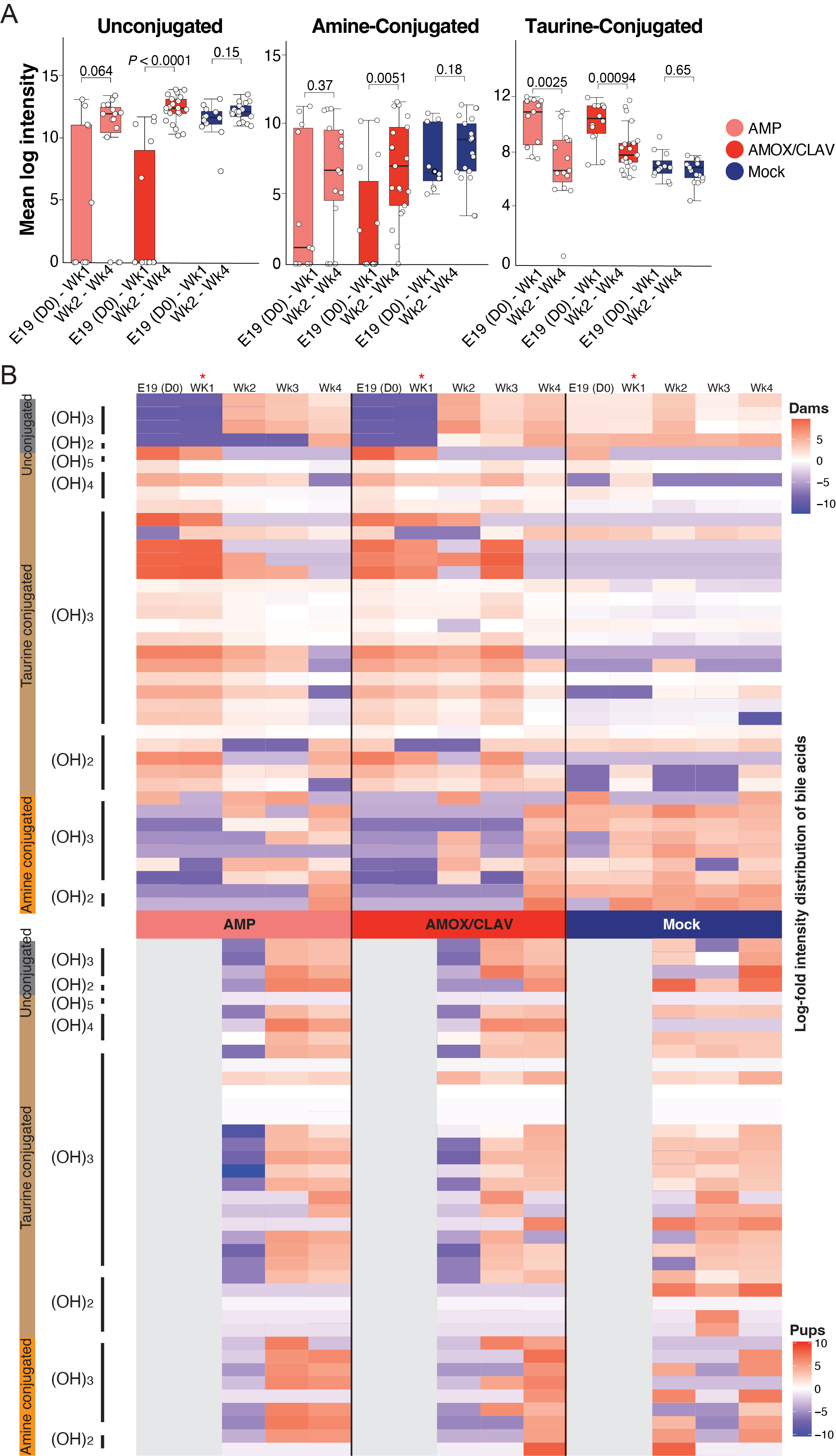

### Supplementary Figure 6

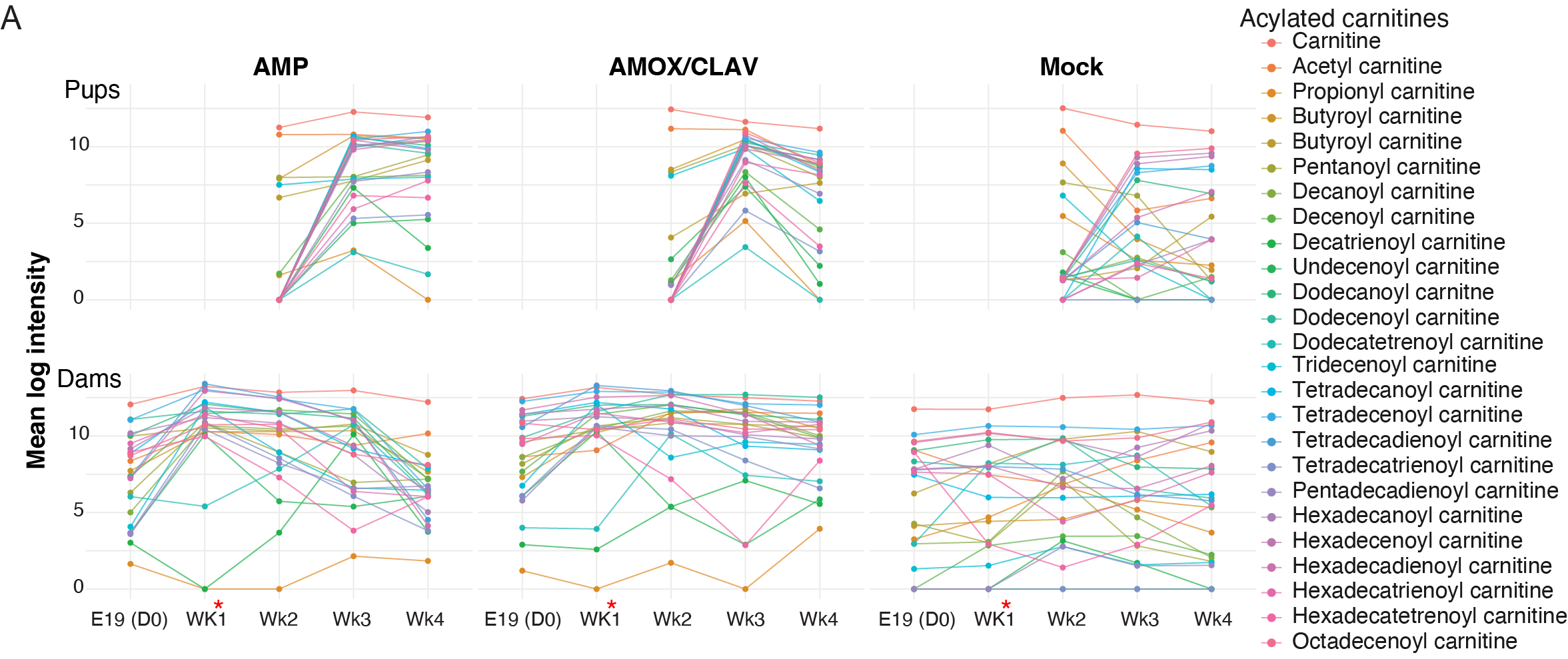

### Supplementary Figure 7

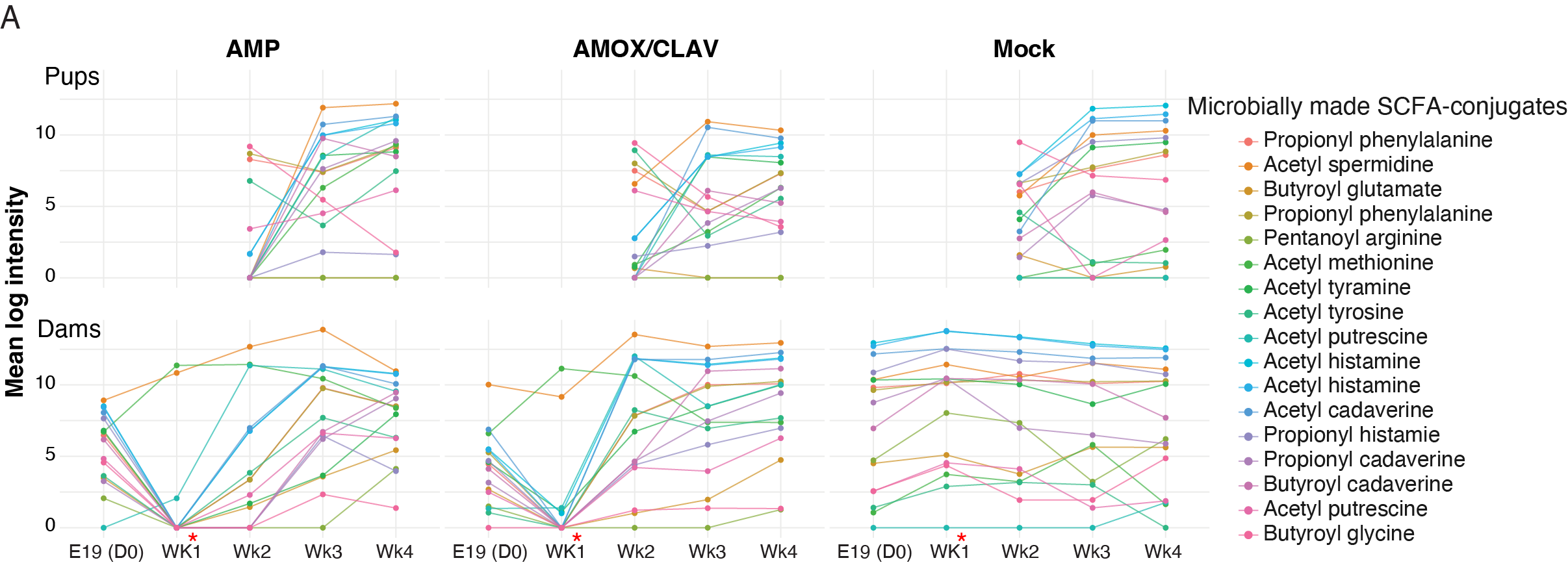
