## Supplementary Information for "Effect of Perinatal Ampicillin or Amoxicillin/Clavulanate Exposure on Maternal and Infant Gut Microbiome, Metabolome, and Infant Responses to the 20-valent Pneumococcal Conjugate Vaccine"

**This file includes:**
Supplementary Figures (including figure legends) S1–S7
Description of Additional Supplementary Files (Dataset Legends)

**Additional supporting materials for this manuscript include the following:**
Datasets S1–S4

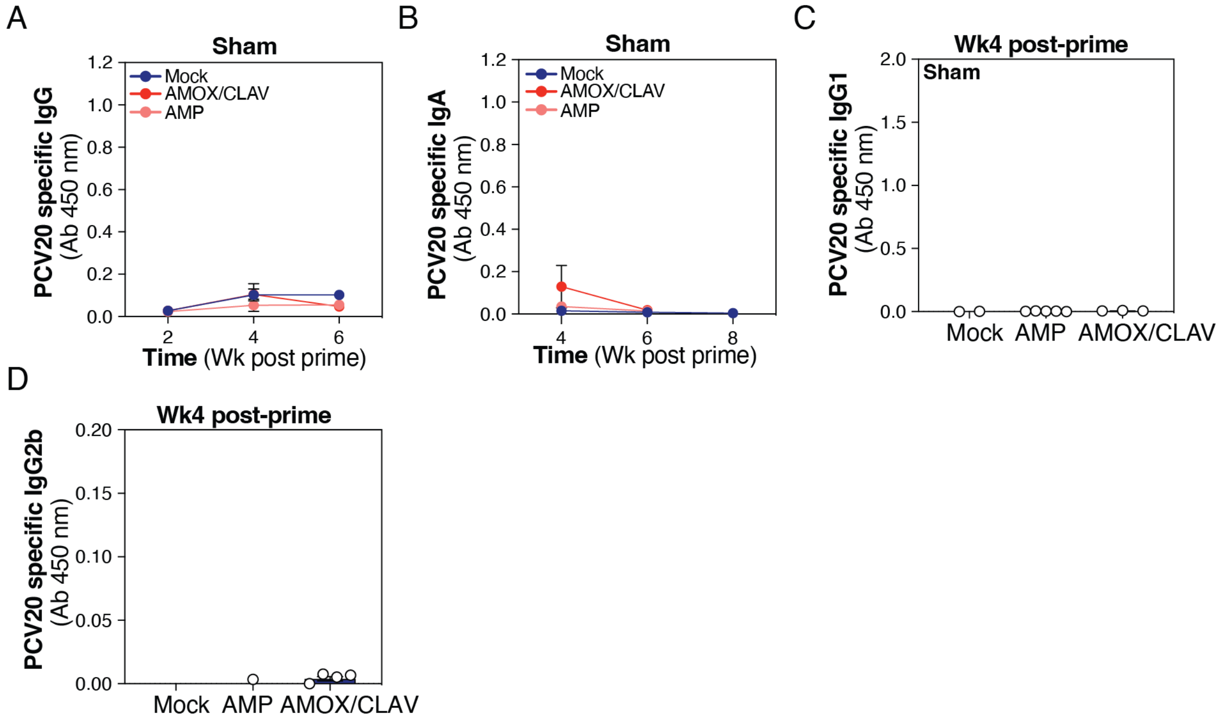


**Figure S1. Peripartum exposure of dams to AMOX/CLAV and their immunological response to PCV20 immunization. (A)** Total serum anti-PCV20 IgG levels were measured following sham (PBS) immunization at the dose of 100 µL/mouse. Serum samples were collected at weeks 2, 4, and 6 post-prime as described in Figure 1A. The PCV20-immunized mice are shown in Fig. 1B. Data are plotted as the mean ± SEM, representing 5-6 mice per group. **(B)** Total serum anti-PCV20 IgA levels were measured following sham (PBS) immunization as described in A. Serum samples were collected at weeks 4, 6, and 8 (mock) post-prime. The PCV20-immunized mice are shown in Fig. 1C. Data are plotted as the mean ± SEM, representing 5-6 mice per group. **(C)** Total serum anti-PCV20 IgG1 levels were measured following sham (PBS) immunization as described in A. Serum samples were collected at week 4 post-prime. The PCV20-immunized mice are shown in Fig. 1D. Data are plotted as the mean ± SEM, representing 5-6 mice per group. **(D)** Total serum anti-PCV20 IgG2b levels were measured following sham (PBS) immunization as described in A. Serum samples were collected at week 4 post-prime. The PCV20-immunized mice are shown in Fig. 1E. Data are plotted as the mean ± SEM, representing 5-6 mice per group.

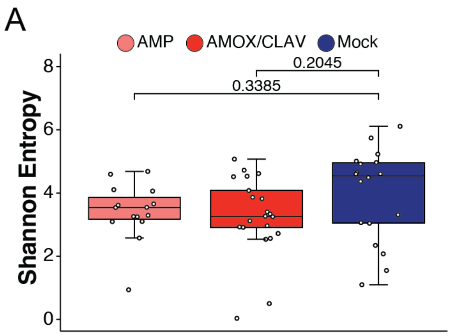


**Figure S2**. **Alpha Diversity analysis of Pups Wk2-4. (A)** Shannon diversity was not significantly ($P$ < 0.05) reduced in pups (regardless of their immunization status) exposed to antibiotic treatments. Statistical significance was assessed using a linear mixed-effects model for repeated measures. The boxplots represent first (lower), interquartile range (IQR), and third (upper) quartile.

A

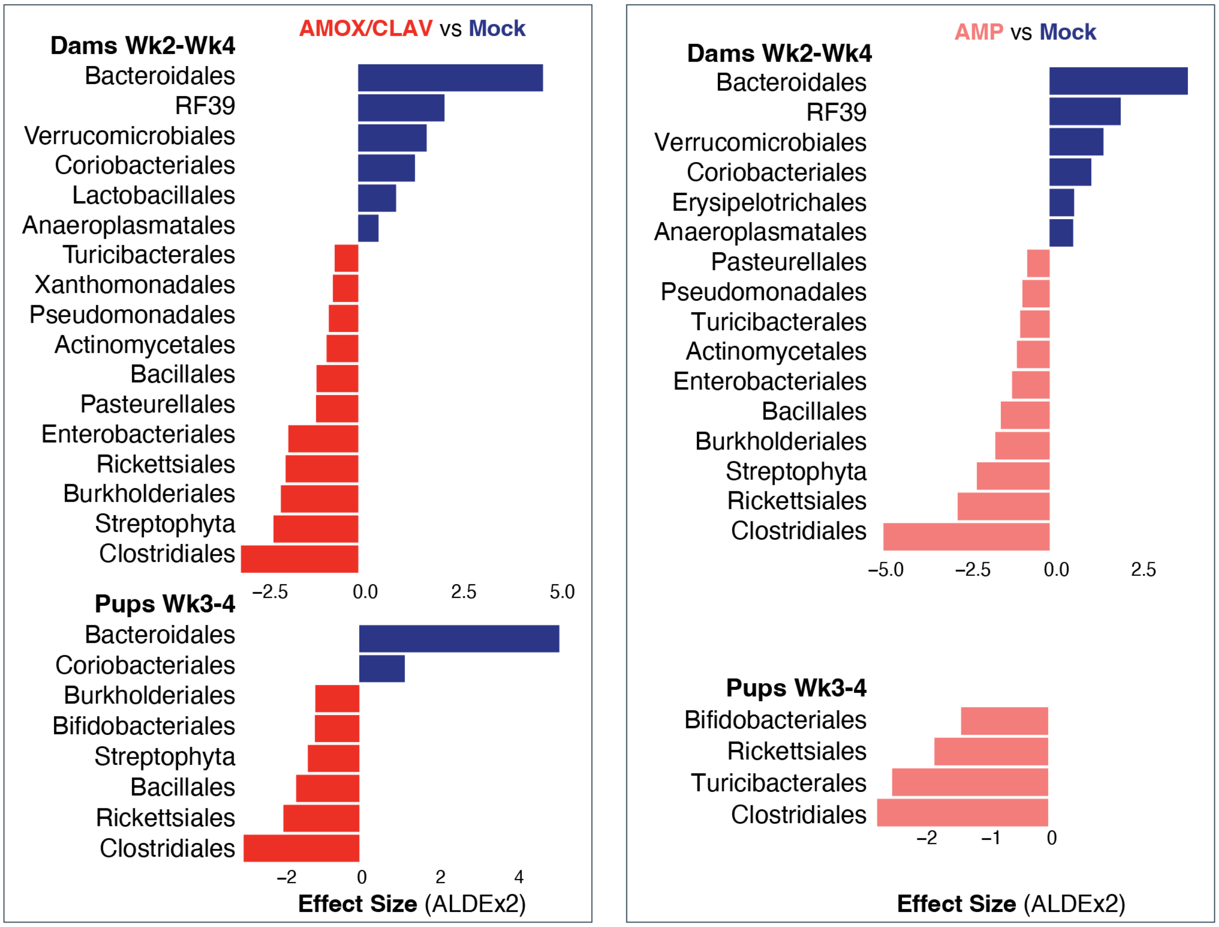


**Figure S3. Microbiome analysis of mice stratified by treatment condition (A)** Key microbial features driving differences between antibiotic treatment conditions at later time points (Dams Wk2-Wk4, PCV-20 immunized pups Wk3-Wk4), collapsed at the order level, were identified using differential abundance analysis with ALDEx2 (FDR-corrected $P < 0.05$).

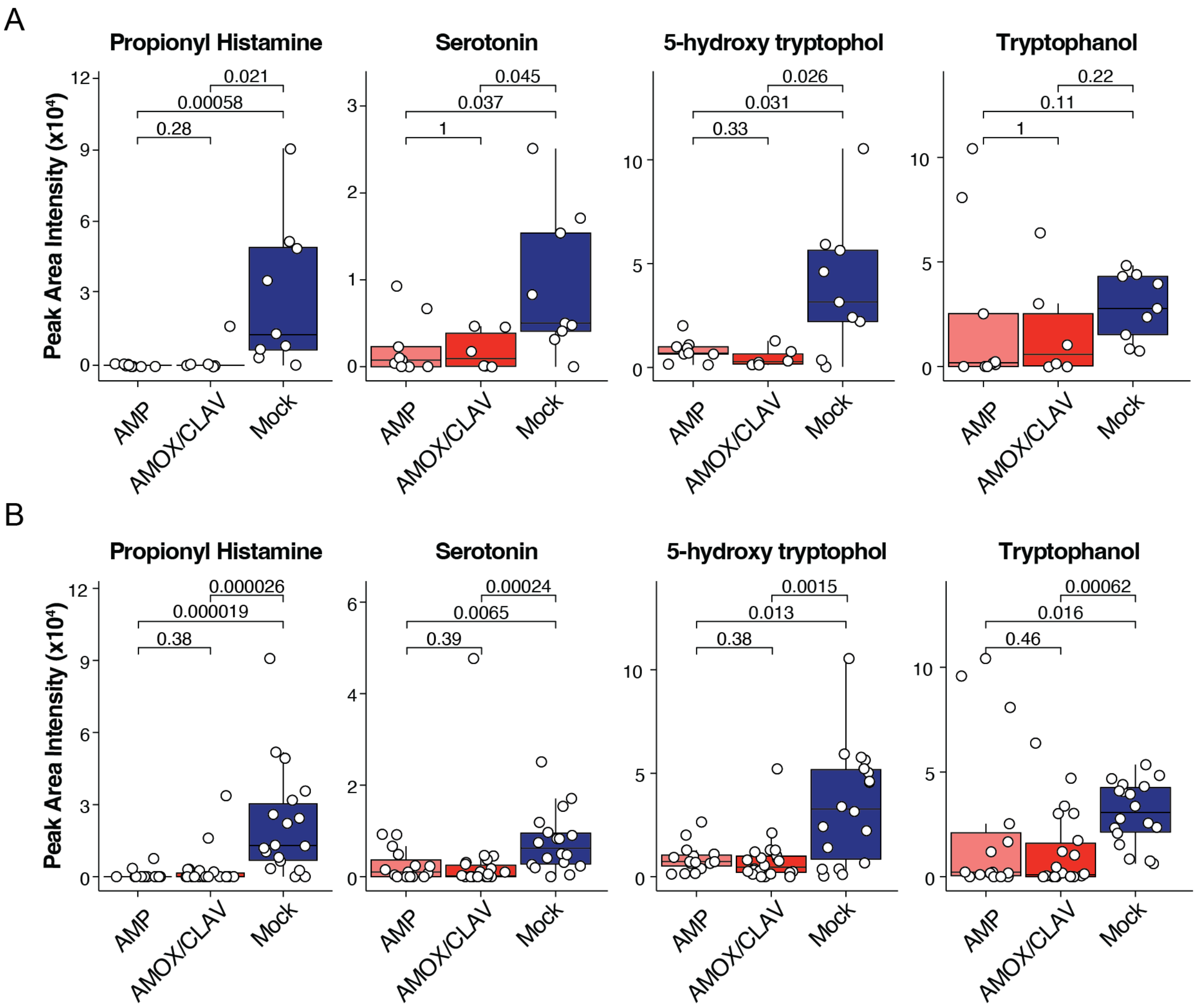


**Figure S4. Tryptophan metabolites across treatment conditions in Pups. (A)** Univariate analysis was performed on selected molecular features: 5-hydroxy tryptophol (ID 808), serotonin (ID 805), propionyl histamine (ID 481), and tryptophanol (ID 2566). Metabolite levels were compared in pups at Wk2 (pre-sham immunization) through WK4 (post-sham immunization) that were indirectly exposed to antibiotics. **(B)** Univariate analysis was performed on selected molecular features: 5-hydroxy tryptophol (ID 808), serotonin (ID 805), propionyl histamine (ID 481), and tryptophanol (ID 2566). Metabolite levels were compared in pups at Wk2 through WK4 (regardless of immunizations status) that were indirectly exposed to antibiotics. Statistical significance (A and B) was assessed using the Wilcoxon rank-sum test with Benjamin-Hochberg false discovery rate correction. The boxplots represent first quartile (lower), interquartile range (IQR), and third (upper) quartile.

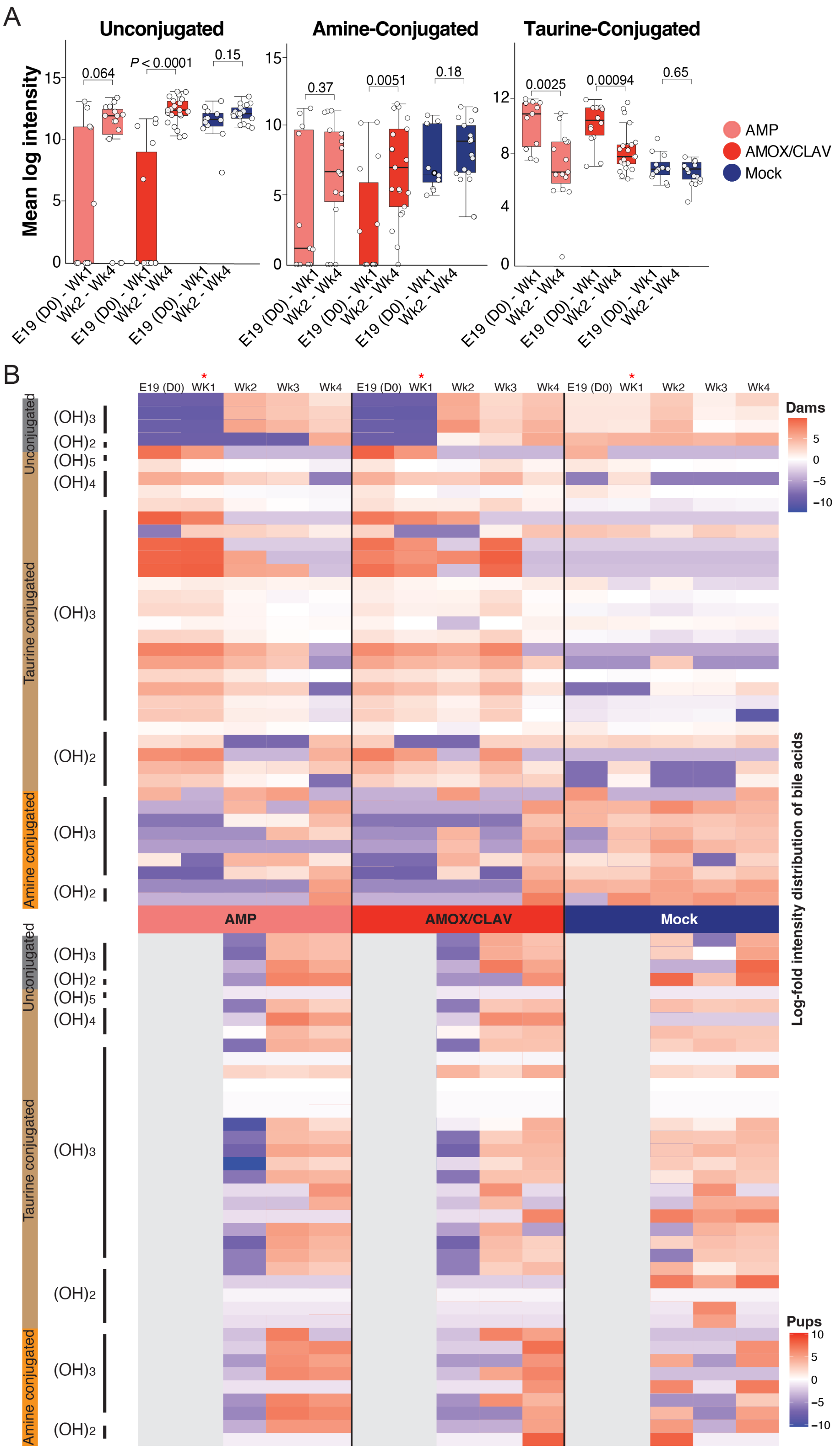

A
Unconjugated
Amine-Conjugated
Taurine-Conjugated
Mean log intensity
0.064
P < 0.0001
0.15
0.37
0.0051
0.18
0.0025
0.00094
0.65
E19 (D0) - Wk1
Wk2 - Wk4
AMP
AMOX/CLAV
Mock
B
E19 (D0)
WK1
Wk2
Wk3
Wk4
Dams
Pups
Unconjugated
Taurine conjugated
Amine conjugated
(OH)3
(OH)2
(OH)5
(OH)4
Log-fold intensity distribution of bile acids

**Figure S5**. **Treatment-specific alterations in bile acid profiles reveal shifts in host and microbial bile acid metabolism. (A)** Box plots comparing mean log intensities of unconjugated, amine-conjugated, and taurine-conjugated bile acids in dams at early (Day 0 to Week 1 postpartum) and late timepoints (Week 2 to Week 4 postpartum). Feature annotations were obtained from the CMMC-enrichment workflow using GNPS2. **(B)** Heat map of annotated bile acid features showing log-fold intensities across treatment groups in dams and pups (pre- and post-immunized with sham or PCV20. Bile acids are categorized by conjugation type: non-conjugated (grey; microbial-derived), amine-conjugated (orange; microbial-derived), and taurine-conjugated (brown; host-derived) spanning diverse hydroxy core structures. Feature annotations were obtained from the CMMC-enrichment workflow using GNPS2. An asterisk indicates the termination of antibiotic treatment.

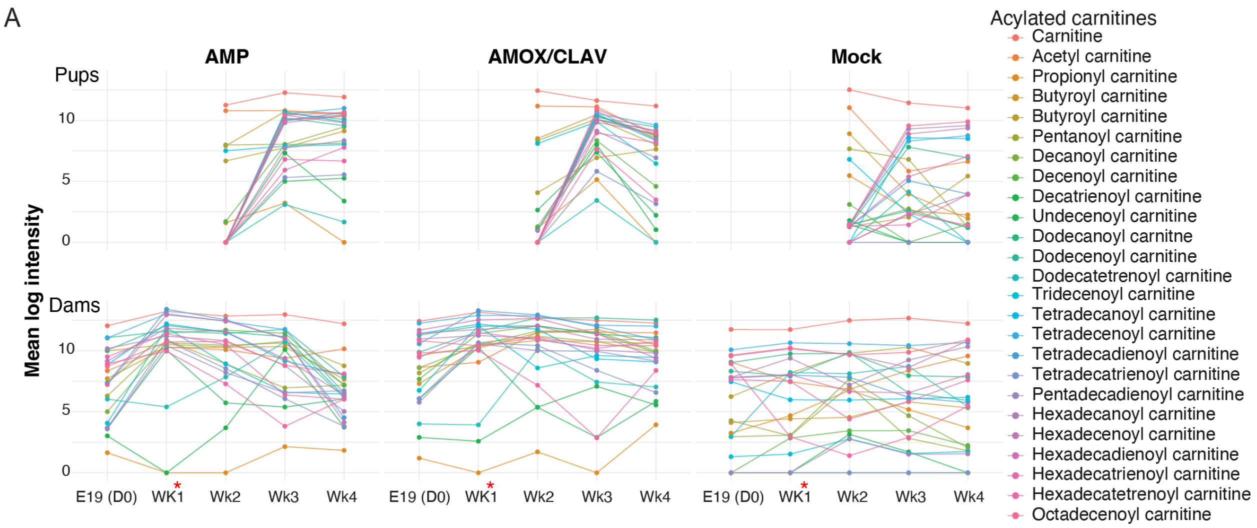


**Figure S6**. **Acyl carnitines dynamics across treatment conditions. (A)** Line graph depicting log intensities of MSI level 2 annotated acyl carnitines, a class of host-associated metabolites, across treatment conditions for dams and pups (pre- and post-immunized with sham or PCV20). Feature annotations were obtained from the CMMC-enrichment workflow using GNPS2. An asterisk indicates the termination of antibiotic treatment.

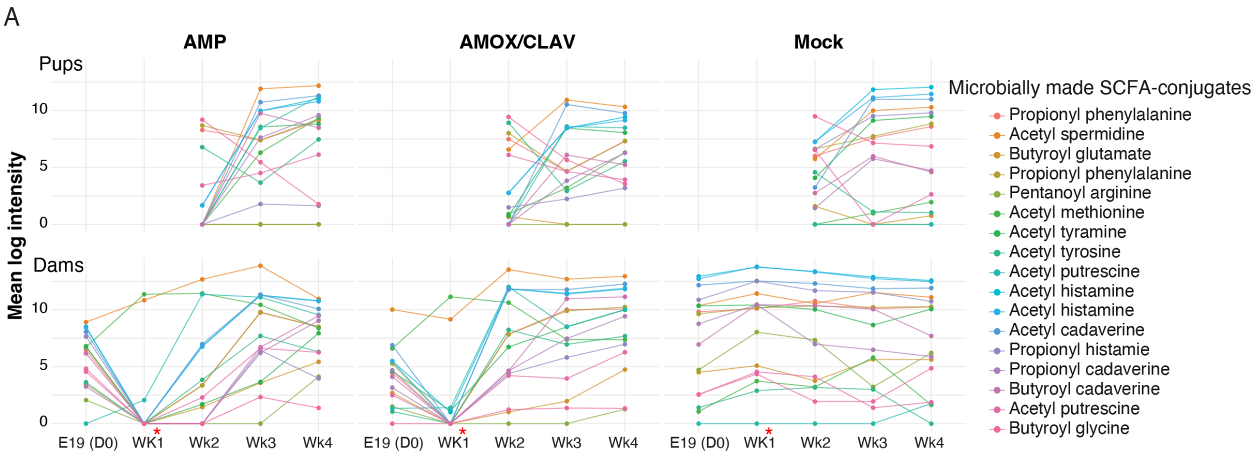


**Figure S7**. ***N*-acyl lipids dynamics across treatment conditions. (A)** Line graph depicting log-fold intensities of annotated *N*-acyl lipids, a class of microbe-associated metabolites, across treatment conditions for dams and pups (pre- and post-immunized with sham or PCV20). Feature annotations were obtained from the CMMC-enrichment workflow using GNPS2. An asterisk indicates the termination of antibiotic treatment.

**Dataset Legend**

**Dataset 1.** Differential abundance analysis of bacterial taxa in dams, stratified by time points and treatment, was performed using ALDEx2, with comparisons made against mock-treated controls. Reported statistical metrics include Welch's t-test *P*-value (we.ep) and Benjamini–Hochberg–adjusted *P*-value (we.eBH), Wilcoxon rank-test *P*-value (wi.ep) and adjusted *P*-value (wi.eBH), centered log-ratio (clr)–transformed relative abundances across all samples (rab.all), within each treatment group (AMP: rab.win.mom.Amp; AMOX/CLAV: rab.win.mom.Aug) and mock-controls (rab.win.mom.Mock), the median clr difference between treatment and controls (diff.btw), within-group variance (diff.win), estimated effect size (effect), and the degree of overlap between posterior distributions (overlap).

**Dataset 2.** Differential abundance analysis of bacterial taxa in pups, stratified by time points and treatment, was performed using ALDEx2, with comparisons made against mock-treated controls. Reported statistical metrics include Welch's t-test *P*-value (we.ep) and Benjamini–Hochberg–adjusted *P*-value (we.eBH), Wilcoxon rank-test *P*-value (wi.ep) and adjusted *P*-value (wi.eBH), centered log-ratio (clr)–transformed relative abundances across all samples (rab.all), within each treatment group (AMP: rab.win.mom.Amp; AMOX/CLAV: rab.win.mom.Aug) and mock-controls (rab.win.mom.Mock), the median clr difference between treatment and controls (diff.btw), within-group variance (diff.win), estimated effect size (effect), and the degree of overlap between posterior distributions (overlap).

**Dataset 3.** Features ranked by Variable Importance in Projection (VIP) scores calculated from the PLS-DA model for component 1. Only features with VIP scores greater than 1 were included and considered important for discriminating between treatment groups. Additional feature details include the PLS-DA loading, mass-to-charge-ratio (mz), retention time (RT), correlation ID (Corr_ID), and compound name. Compound name annotations were obtained using the feature based molecular networking workflow on GNPS2.

**Dataset 4.** Comparison of LC–MS/MS extracted ion feature abundances in serum of pups (Wk1 postpartum) revealed no detectable features corresponding to either AMOX/CLAV or AMP. A 10 µM pure amoxicillin was included as standard and injected at the end of the sequence.
